# The neuronal fate determinants SOX4/11 control mitotic fidelity of adult hippocampal precursor cells

**DOI:** 10.1101/2025.10.13.682016

**Authors:** Sneha Adhikarla, Iris Schäffner, Angélica Luna Leal, Shailendra Kumar Singh, Benjamin Martin Häberle, Sören Turan, Elizabeth Sock, Andreas Sagner, Dieter Chichung Lie

**Affiliations:** Institute for Anatomy, Friedrich-Alexander Universität Erlangen-Nürnberg, 91054 Erlangen, Germany; Institute of Biochemistry, Friedrich-Alexander Universität Erlangen-Nürnberg, 91054 Erlangen, Germany

**Keywords:** mitosis, cell death, neural precursor cells, neurogenesis, SOXC transcription factors

## Abstract

In adult hippocampal neurogenesis, fast dividing intermediate progenitor cells (IPCs) ensure the production of a larger number of neurons from a limited pool of slow dividing radial-glia-like neural stem cells. Here, we demonstrate that the neuronal fate determining and lineage-specific transcription factors SOX4 and SOX11 are essential to faithfully execute mitosis in IPCs. In vivo, combined deletion of SOX4 and SOX11 results in death of IPCs and abolishes the generation of new neurons. In vitro analyses of SOX4/11-deficient precursors revealed mitosis defects including chromosome segregation errors, centrosomal errors and cytokinesis defects. SOX4/11-deficient precursors frequently featured micronuclei and DNA bridges and showed a pro-inflammatory signaling profile, suggesting the induction of death by mitotic catastrophe. Importantly, analysis of the developing mouse spinal cord and of human pluripotent stem cell-derived brain organoids indicate that SOXC transcription factors are essential for mitotic fidelity of neural precursor cells across ontogeny and species. The data raise the interesting possibility that mitotic programs in precursor cells are controlled in a lineage-specific manner.

## Introduction

The dentate gyrus of the adult hippocampus is a dynamic structure in which new neurons are generated throughout life in many mammalian species including humans (Dumitru *et al*, 2025; Eriksson *et al*, 1998; Goncalves *et al*, 2016; Kuhn *et al*, 2018; Moreno-Jimenez *et al*, 2019; Spalding *et al*, 2013; Terreros-Roncal *et al*, 2021; Zhou *et al*, 2022) The continuous generation of dentate granule neurons, which is also termed adult hippocampal neurogenesis, is essential for hippocampus-dependent plasticity; whereas impaired adult neurogenesis has been suggested to contribute to cognitive and behavioral symptoms in neurodegenerative and neuropsychiatric disorders and ageing (Ahlenius *et al*, 2009; Boldrini *et al*, 2018; Demars *et al*, 2010; Hill *et al*, 2015; Tobin *et al*, 2019; Unger *et al*, 2016). Adult-born dentate granule neurons are generated from radial glia like stem cells (RGLs) through a multistep process, involving activation of the quiescent RGL, generation of fast dividing intermediate precursor cells (IPCs), neuronal fate determination and maturation (Kempermann *et al*, 2015). IPCs undergo two to three rounds of rapid division to expand the precursor pool (Pilz *et al*, 2018) thereby enabling the generation of a larger number of new neurons from a limited stem cell pool. Expansion of the neuronal precursor pool through IPC proliferation is critical, considering the negative impact of decreased adult dentate granule neuron production on hippocampus-dependent plasticity (Abrous & Wojtowicz, 2015; Tuncdemir *et al*, 2019). Rapid cell division, however, is thought to render cells such as IPCs vulnerable to mitosis errors (Levine & Holland, 2018). Indeed, in the developing mouse brain about a third of rapidly cycling neural precursor cells were reported to feature mitosis defects such as chromosome segregation errors or presence of multiple centrosomes (Rehen *et al*, 2001; Yang *et al*, 2003). Irreparable and deleterious mitotic defects in turn have been suggested to trigger cell death of precursor cells in order to prevent the generation and integration of functionally aberrant neurons into brain circuits (McKinnon, 2013). Whether neural progenitor cells entertain specialized mechanisms to safeguard mitotic fidelity, however, is largely unknown.

SOX4 and SOX11 are members of the SOXC family of transcription factors (Wegner, 2011). They display a highly overlapping expression pattern in the developing and adult central nervous system that is restricted to the neuronal lineage. SOX4 and SOX11 fulfill pleiotropic functions in neurogenesis such as the regulation of neuronal fate determination, neuronal migration, dendrite growth, and timing of synaptic integration (Bergsland *et al*, 2006; Chen *et al*, 2015; Hoshiba *et al*, 2016; Mu *et al*, 2012; Wang *et al*, 2013). SOX4 and SOX11 also function redundantly in neural precursor survival in embryonic mouse development (Bhattaram *et al*, 2010; Thein *et al*, 2010) and it has been suggested that part of this phenotype is the consequence of dysregulated Hippo signaling (Bhattaram *et al*., 2010). Here, we uncover that neuronal fate determinants of the SOXC family are essential for survival of IPCs in adult neurogenesis by safeguarding mitotic fidelity. We also demonstrate that SOXC factors are important for mitotic fidelity of precursor cells across ontogeny and species, thereby raising the novel concept that neuronal lineage-specific mitotic programs are wired into neuronal fate determining programs.

## Results

### SOX4 and SOX11 are expressed in adult neural stem and progenitor cells

We first analyzed the expression of SOX4 and SOX11 in transgenic mice, in which eGFP is expressed under the control of the RGL-specific NESTIN-promoter (NESTIN::eGFP mice). Activated RGLs were distinguished from quiescent RGLs by the co-expression of the proliferation marker MCM2, while IPCs were identified using the IPC-specific transcription factor TBR2. Consistent with transcriptomic analyses (Rasetto *et al*, 2024; Shin *et al*, 2015), we found that only a small subset of quiescent RGLs expressed SOX4 and SOX11 (13% and 8%, respectively, Figure 1A, C). SOX4 and SOX11 expression strongly increased in activated RGLs (100% and 63%, respectively, Figure 1A, C), and virtually all TBR2^+^ IPCs expressed both SOX4 and SOX11 (100% and 91%, respectively) (Figure 1B, C).

**Figure 1:**
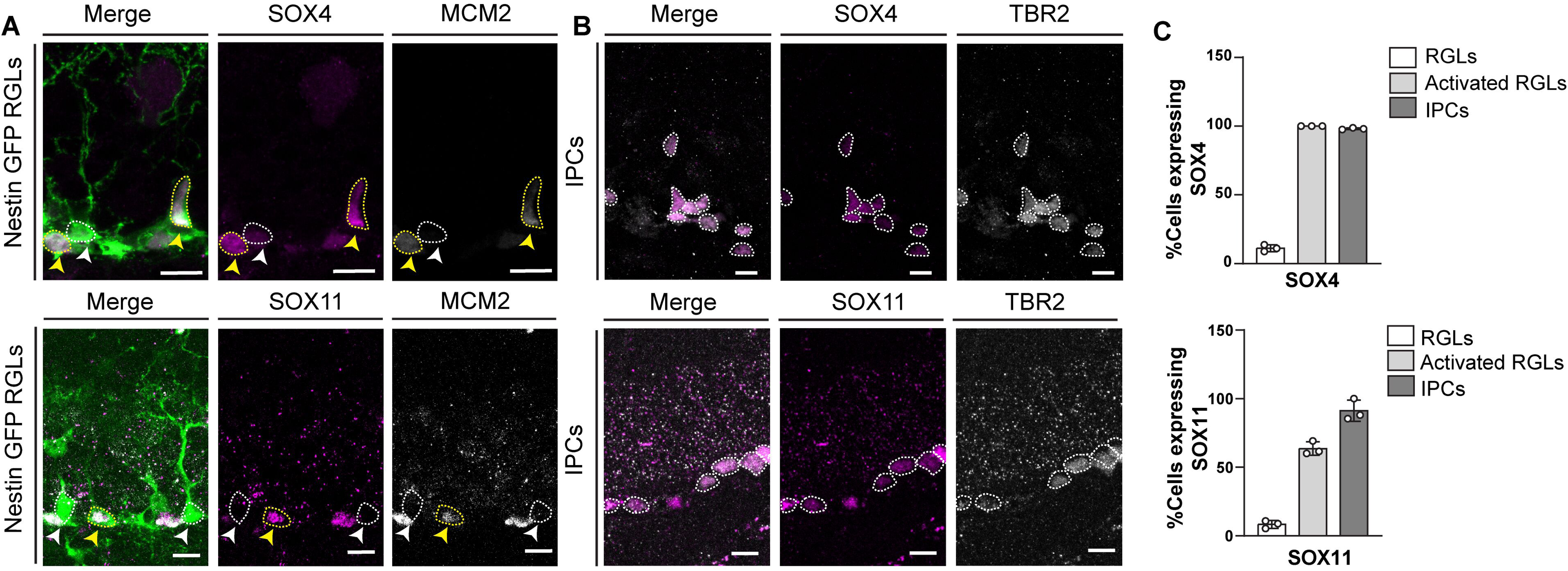
SOX4 and SOX11 are strongly expressed in activated RGLs and IPCs. **(A)** Representative images showing co-localization of SOX4 (magenta upper panel) and SOX11 (magenta lower panel) with MCM2 (grey) and Nestin-GFP (green) in activated RGLs (yellow arrowheads). MCM2-quiescent RGLs (white arrowheads). Scale bar = 10µm. **(B)** Representative images showing expression of SOX4 (magenta upper panel) and SOX11 (magenta lower panel) in TBR2^+^ (grey) IPCs. Scale bar= 10µm. **(C)** Quantification of percentage of each cell type expressing SOX11 (upper graph) and SOX4 (Lower graph). 50 to 100 cells per animal analyzed, n = 3. ***P<0.001, **P<0.01, *P<0.05, ns = non-significant; dots represent individual animals

### SOXC deficiency impairs generation of immature neurons in the adult dentate gyrus

To investigate the function of SOXC proteins in early developmental stages of adult neurogenesis, we generated RGL-specific SOX4/SOX11 conditional knockout mice (Nestin::CreERT2; Rosa26floxSTOPfloxYFP; Sox4flox/flox Sox11flox/flox mice; hereafter referred to as SOX4/11*^iNestin^*), which allow for tamoxifen-induced deletion of SOX4 and SOX11 in RGLs and tracing of recombined cells by expression of YFP. Nestin::CreERT2; Rosa26floxSTOPfloxYFP mice (hereafter referred to as control) harboring wildtype alleles for SOX4 and SOX11 served as control. Recombination was induced in young adult (8-10 week old) mice (Figure 2A).

**Figure 2:**
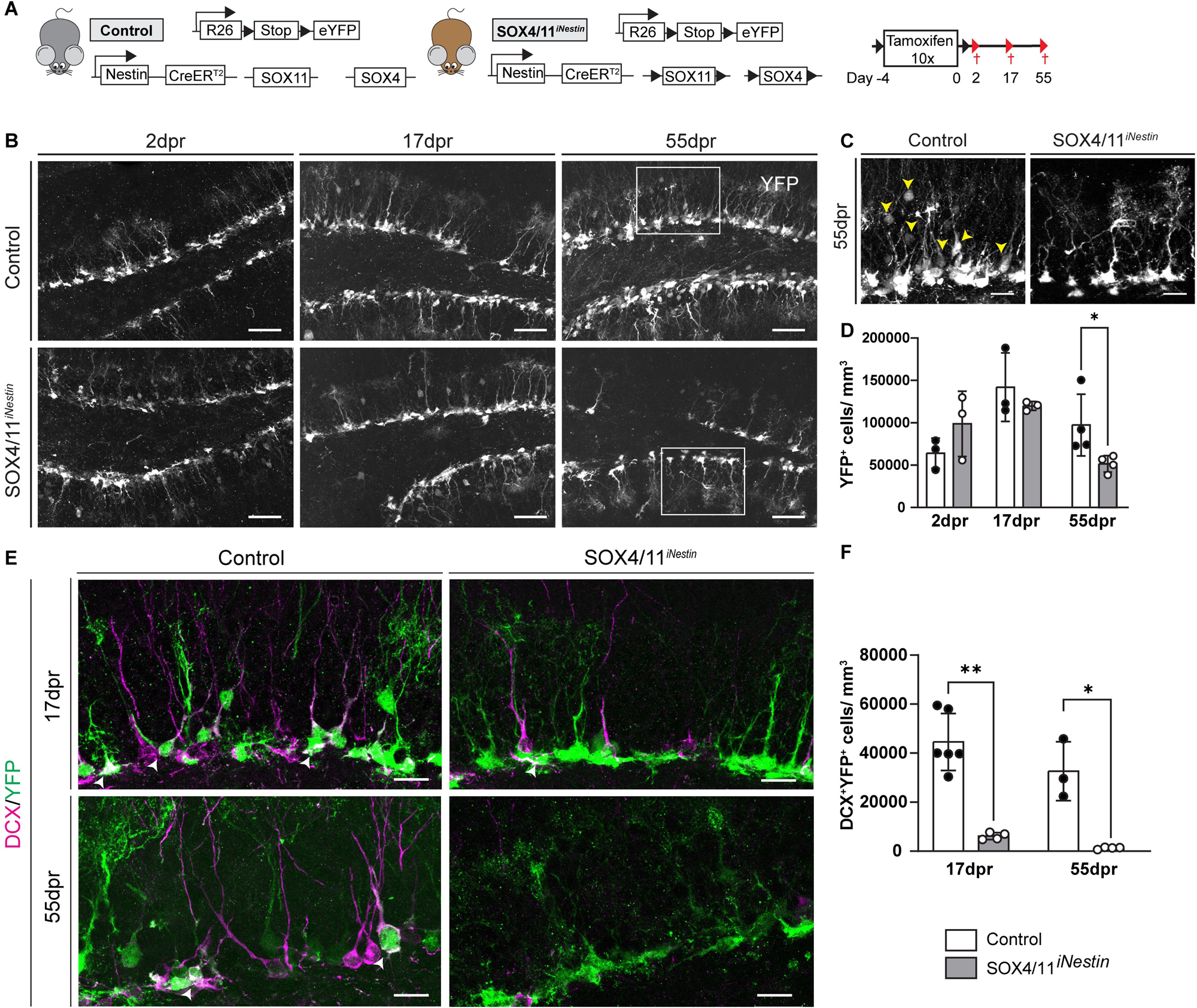
SOX4/11 deletion ablates the generation of new neurons. **(A)** Schematic representation of the conditional alleles and transgenes and experimental scheme used in (B) –(E). **(B)** Representative images showing YFP^+^ cells in the dentate gyrus at 2dpr (left), 17dpr (middle) and 55dpr (right). Scalebar = 50µm. **(C)** Zoomed in panel from 55dpr showing YFP^+^ cells with mature neuron morphology (yellow arrows) and RGL like morphology in Control mice (left panel) while YFP^+^ cells in SOX4/11^iNestin^ mice show only a RGL like morphology (right panel). **(D)** Quantification of YFP^+^ cells in the sub-granular zone of the dentate gyrus. Note the loss of YFP^+^cells with neuronal morphology on day 55 post recombination in SOX4/11^iNestin^ mice. n= 3-4 animals. **(E)** Representative images of the dentate gyrus of control and SOX4/11^iNestin^ mice at 17dpr (left) and 55dpr (right). Note the loss of DCX^+^ (magenta) YFP^+^ (green) immature neurons (white arrowheads) in SOX4/11^iNestin^ mice at 17dpr (left) and 55dpr (right). Scalebar= 20µm. **(F)** Quantification of YFP^+^DCX^+^ cells at 17dpr and 55dpr. n= 3-6 animals. Data represented as mean ± SD; t test was used to determine significance. ***P<0.001, **P<0.01, *P<0.05, ns = non-significant; dots represent individual animals

On day 2 post induction of recombination (2dpr), SOX4/11*^iNestin^*and control mice showed comparable numbers of YFP^+^ cells, indicating that the recombination efficiency was comparable between experimental groups (Figure 2B, C). Moreover, YFP^+^ cells in SOX4/11*^iNestin^* mice were negative for SOX4 and SOX11 demonstrating their successful conditional ablation (Expanded View, EV 2A).

The effects of SOX4/11 deletion on neurogenesis were evaluated at 17 dpr and 55 dpr. While the number of YFP^+^ cells was comparable between control and SOX4/11*^iNestin^* at 17dpr (Figure 2B-D), there was a significant decrease in the number of YFP^+^ cells in SOX4/11*^iNestin^* at 55 dpr. SOX4/11*^iNestin^* mice showed in particular a depletion of YFP^+^ cells with neuronal morphology, suggesting a block in the generation of neurons (Figure 2C). To further substantiate the impairment in generation of new dentate granule (DG) neurons, sections were stained for the immature neuronal markers doublecortin (DCX) (Figure 2E) and NEUROD1 (EV Figure 2B), and for the DG neuron marker PROX1 (EV Figure 2C). As expected, numerous recombined cells (YFP^+^) expressing neuronal markers were found in control mice at 17dpr. However, YFP^+^ cells bearing neuronal markers were almost completely absent in SOX4/11*^iNestin^* mice (Figure 2E, F; EV Figure 2 B-E), indicating that SOX4/11 deficiency abrogated or delayed the formation of new DG neurons. At 55 dpr, the dentate gyrus of control mice harbored numerous recombined DCX^+^ neurons, while YFP^+^ immature neurons remained virtually absent in SOX4/11*^iNestin^* mice (Figure 2E, F), demonstrating that SOX4/11 deficiency in the adult neurogenic lineage impaired rather than delayed the generation of DG neurons.

### Deletion of SOXC transcription factors results in cell death of IPCs

We next examined at which developmental stage SOX4/11 deficiency impaired neurogenesis. At 17dpr, SOX4/11*^iNestin^* mice showed increased numbers of recombined RGLs, and an increase in activated RGLs (EV Figure 3A-C) and TBR2^+^ IPCs (Figure 3A, B) respectively, compared to the control mice. These data indicate that SOX4/11 deficiency neither impaired RGL activation nor ablated the differentiation of RGLs into IPCs. Notably, the number of YFP^+^ IPCs dropped between 17 dpr and 55 dpr (Figure 3A, B) excluding the possibility that SOX4/11-deficient cells arrested and accumulated at the IPC stage.

**Figure 3:**
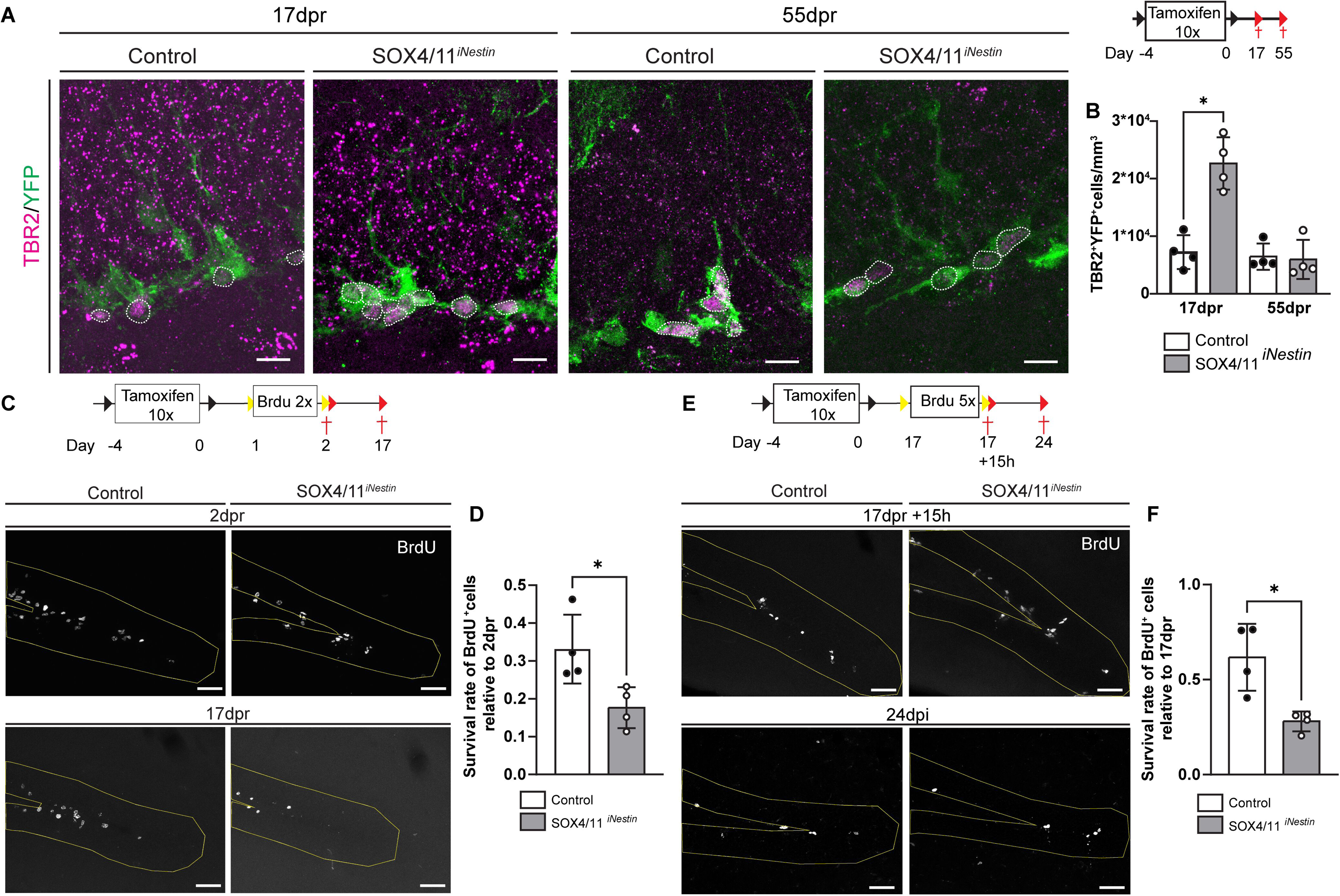
SOX4/11 are required for IPC survival during adult neurogenesis. **(A)** Experimental paradigm and representative images showing TBR2^+^ (magenta) recombined cells (green) in the dentate gyrus at 17dpr (left) and 55dpr (right). Scale bar= 20µm **(B)** Quantification forTBR2^+^ YFP^+^ cells at 17 and 55 dpr in the subgranular zone of the dentate gyrus, showing transient increase of progenitors at 17dpr in SOX4/11^iNestin^ mice. n=4 animals. Experimental timeline and representative images of BrdU^+^ cells in the dentate gyrus (outlined in yellow) at 2dpr and 17dpr. Scale bar= 50µm **(D)** Survival rate calculated as the number of BrdU^+^ cells at 17dpr relative to the number of of BrdU^+^ cells at 2dpr. SOX4/11^iNestin^ mice show a decreased survival rate of the BrdU^+^ cells. n=4 animals. **(E)** Experimental timeline and representative images of BrdU^+^ cells in the dentate gyrus (outlined in yellow) at 17dpr +15hrs and at 24 dpr. Scale bar= 50µm. **(F)** Survival rate calculated as the number of BrdU^+^ cells at at 24 dpr relative to the number of of BrdU^+^ cells at 17dpr +15hrs. SOX4/11^iNestin^ mice show a decreased survival rate of the BrdU^+^ cells. n= 4 animals. Data represented as mean ± SD; t test was used to determine significance. ***P<0.001, **P<0.01, *P<0.05, ns = non-significant; dots represent individual animals

We next determined whether IPCs gave rise to glia instead of neurons and analyzed expression of the pan-oligodendrocytic marker OLIG2 and the pan-astrocytic marker S100B in recombined cells. No YFP^+^ oligodendrocytes were observed in control or SOX4/11*^iNestin^* mice and we found no substantial increase in the number of YFP^+^ S100β^+^ astrocytes in SOX4/11*^iNestin^*mice (EV Figure 3D, E). Collectively, these data exclude the possibility that IPCs switched fate and entered the glial lineage.

Next, we sought to analyze cell death in the neurogenic lineage. Cells positive for the apoptosis marker cleaved caspase 3 (CC3) were in low abundance and highly variable in number, hence rendering the data inconclusive. We therefore opted to estimate survival of newborn cells using BrdU pulse-chasing. Mice were injected with BrdU for 2 days after the final tamoxifen injection and analyzed either 3 hrs (i.e. 2 dpr) or 15 days (i.e. 17 dpr) after the last BrdU injection (Figure 3C). Indeed, the survival rate estimated as the number of BrdU^+^ cells at late time point normalized to the number of BrdU^+^ cells at early time point was substantially decreased in SOX4/11*^iNestin^* (Figure 3D).

As described above, we found a substantial expansion of the highly proliferative IPC pool at 17 dpr in SOX4/11*^iNestin^* mice, which was no longer present at 55dpr (Figure 3A). To estimate the survival of the 17day IPC pool, mice were injected five times with BrdU on 17 dpr and analyzed 3 hrs (i.e., 17dpr+15hrs) or 7 days (i.e., 24dpr) after the last BrdU injection (Figure 3E). Again, SOX4/11*^iNestin^* mice showed a significantly lower survival rate of BrdU^+^ cells compared to controls (Figure 3F), further substantiating the notion that SOX4/11-deficiency impaired survival of IPCs.

### Adult hippocampal neural stem/precursor cells show mitotic apparatus defects upon SOXC deletion

To study the role of SOX4/11 in precursor cell survival in more detail, we established adult neural stem/progenitor cultures (aNSPCs) from the dentate gyrus of young adult mice (8 - 9 weeks old) harboring conditional alleles for SOX4 and SOX11. Cells were transduced with a mouse moloney leukemia retrovirus (MML-V) bicistronically encoding for GFP and Cre-recombinase to generate SOX4/11-deficient aNSPCs (SOX4/11^dKO^ cells). Cells transduced with a MML-V encoding for GFP only served as controls (control cells). aNSPCs were kept in proliferating conditions and collected on day 3 after transduction (Figure 4A). qPCR and Western Blot analyses confirmed downregulation of SOX4 and SOX11 at the transcript as well as the protein level (EV Figure 4A-C). Compared to GFP^+^ control cells, GFP^+^ SOX4/11^dKO^ cells showed slower expansion over a three-day observation period (Figure 4B, C). In addition and in line with the finding that SOX4/11-deficiency impaired precursor survival in vivo, we observed a strong trend towards higher number of apoptotic (cCASP3^+^) cells amongst transduced cells (GFP^+^) in SOX4/11^dKO^ cells compared to control cells (Figure 4D,E).

**Figure 4:**
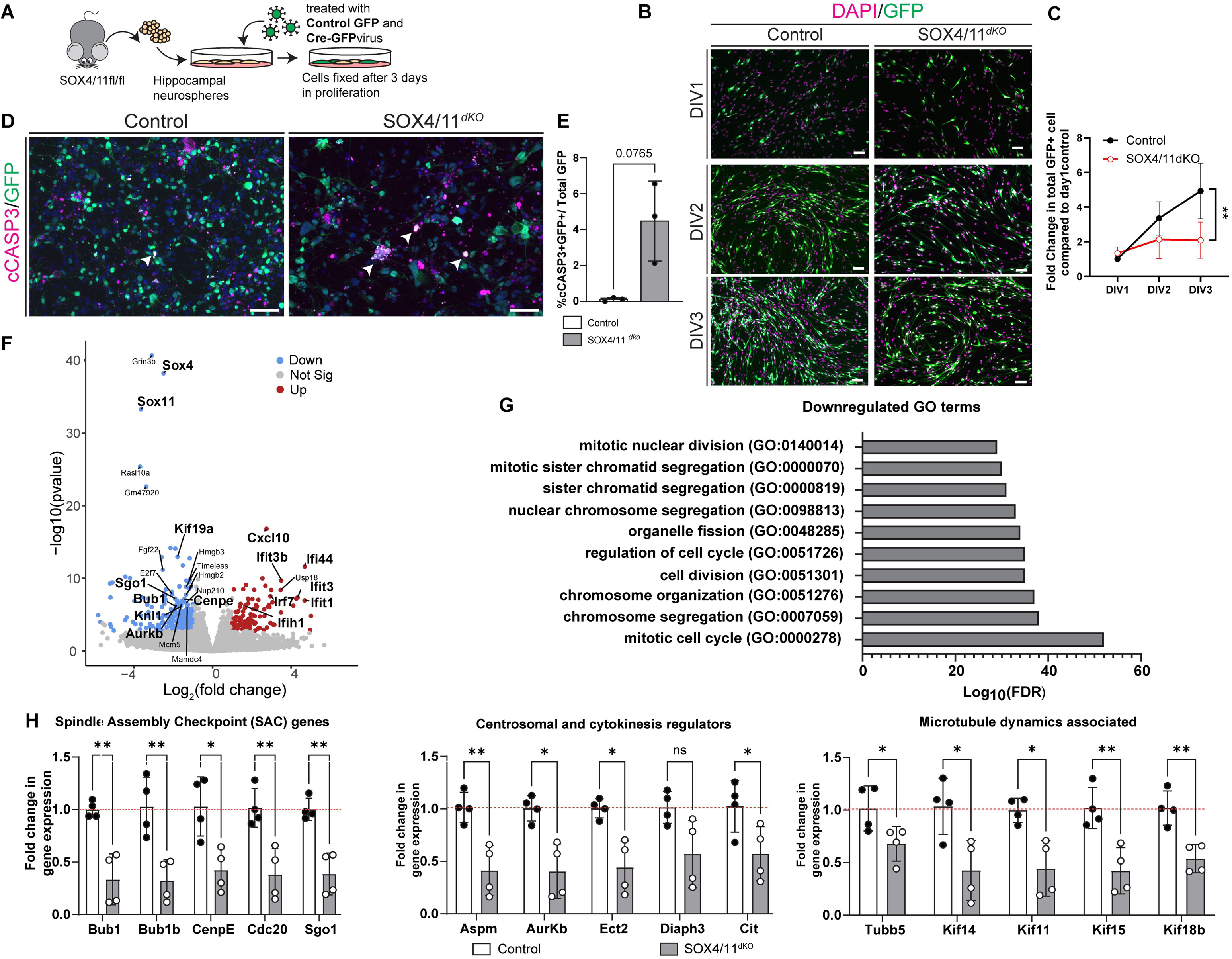
SOX4/11-deficiency in aNSPCs results in increased cell death and transcriptomic defects: **(A)** Experimental scheme used in (B) – (J). **(B)** Representative images of SOX4/11^dKO^ and control cultures on DIV1-3. Virus transduced cells are identified by GFP signal (green), nuclei are counterstained with DAPI (magenta). Scale bar = 50µm. **(C)** Quantification of GFP^+^ cells among all cells at DIV1-3. n=3 biological replicates. **(D)** Representative images of SOX4/11^dKO^ and control cultures stained for cCASP3 (magenta) on DIV3. aNSPCs showed increased cell death as seen in representative image cCASP3 (magenta). Transduced cells are identified by GFP signal (green). Note the increased number of cCASP3^+^ cells. Scale bar = 50µm. **(E)** Quantification of cCASP3^+^GFP^+^ shows a trend towards increased apoptotic cells in SOX4/11^dKO^ cultures, n=3 biological replicates. **(F)** Bulk RNA-Seq experiment was performed at 3 DIV. n= 3 independent biological replicates. Volcano plot showing dysregulated genes. Significantly (fold change=1 and p-value=0.05) up- and downregulated genes are shown in red and blue, respectively. **(G)** GO term analysis of 273 downregulated genes shows enrichment of terms related to mitosis. **(H)** qRT-PCR validation of selected downregulated genes regulating key processes during mitosis such as (left to right) Spindle assembly checkpoint (SAC), Centrosomal and cytokinesis regulators and microtubule dynamics associated genes. n=4 biological replicates. Data represented as mean ± SD; t test was used to determine significance. ***P<0.001, **P<0.01, *P<0.05, ns = non-significant; dots represent individual animals

To gain insight into the molecular impact of SOX4/11 deletion on aNSPCs, we carried out comparative transcriptomic analysis. 382 genes were found to be differentially expressed between SOX4/11^dKO^ and control cells (log_2_fold change >1, adjusted p-value<0.05), among which 109 genes were upregulated and 273 genes were downregulated in SOX4/11^dKO^ cells (Figure 4F). Gene ontology (GO) analysis of differentially expressed genes showed a conspicuous enrichment of genes linked to mitosis and cell-cycle amongst downregulated genes (Figure 4G). qPCR analysis confirmed downregulation of genes encoding for proteins with functions in the mitotic spindle assembly checkpoint (Bub1, Bub1B, CenpE, Cdc20 and Sgo1), spindle formation and chromosome movement (Tubb5, Kif11,Kif14, Kif15 and Kif18b) and the centrosome (ASPM, Cit, ECT2, AurkB, DIAPH3) (Figure 4H), indicating broad perturbation in the expression of genes related to the formation of the mitotic spindle apparatus. To determine whether SOX4/11^dKO^ affected mitotic spindle architecture, we analyzed the parameters mitotic spindle area and length (Kletter *et al*, 2022). Mitotic spindle area in retrovirus transduced cells (i.e., GFP^+^ cells) was quantified by analyzing the immunostaining signals for the spindle microtubules acetylated α-Tubulin (Acetyl α-Tub) and βTubulin5 (Tubb5) (Breuss *et al*, 2016), and the spindle-pole enriched microtubule minus end organizing protein abnormal spindle-like microcephaly-associated (ASPM) (Figure 5A). Morphometric analysis revealed a significant reduction in the area covered by each of these mitotic spindle components in SOX4/11^dKO^ aNSPCs compared to control aNSPCs (Figure 5B). For determination of mitotic spindle length, cells were stained for the centrosome-specific protein γ-tubulin and the distance between the centrosomes was calculated, which was significantly reduced in knockout cells compared to control cells (Figure 5C, D). During this analysis, we also frequently observed supernumerous centrosomes in SOX4/11^dKO^ aNSPCs (EV Figure 5A, B). In sum, these morphometric analyses demonstrate that loss of SOX4/11 caused substantial defects in mitotic spindle architecture of aNSPCs.

**Figure 5:**
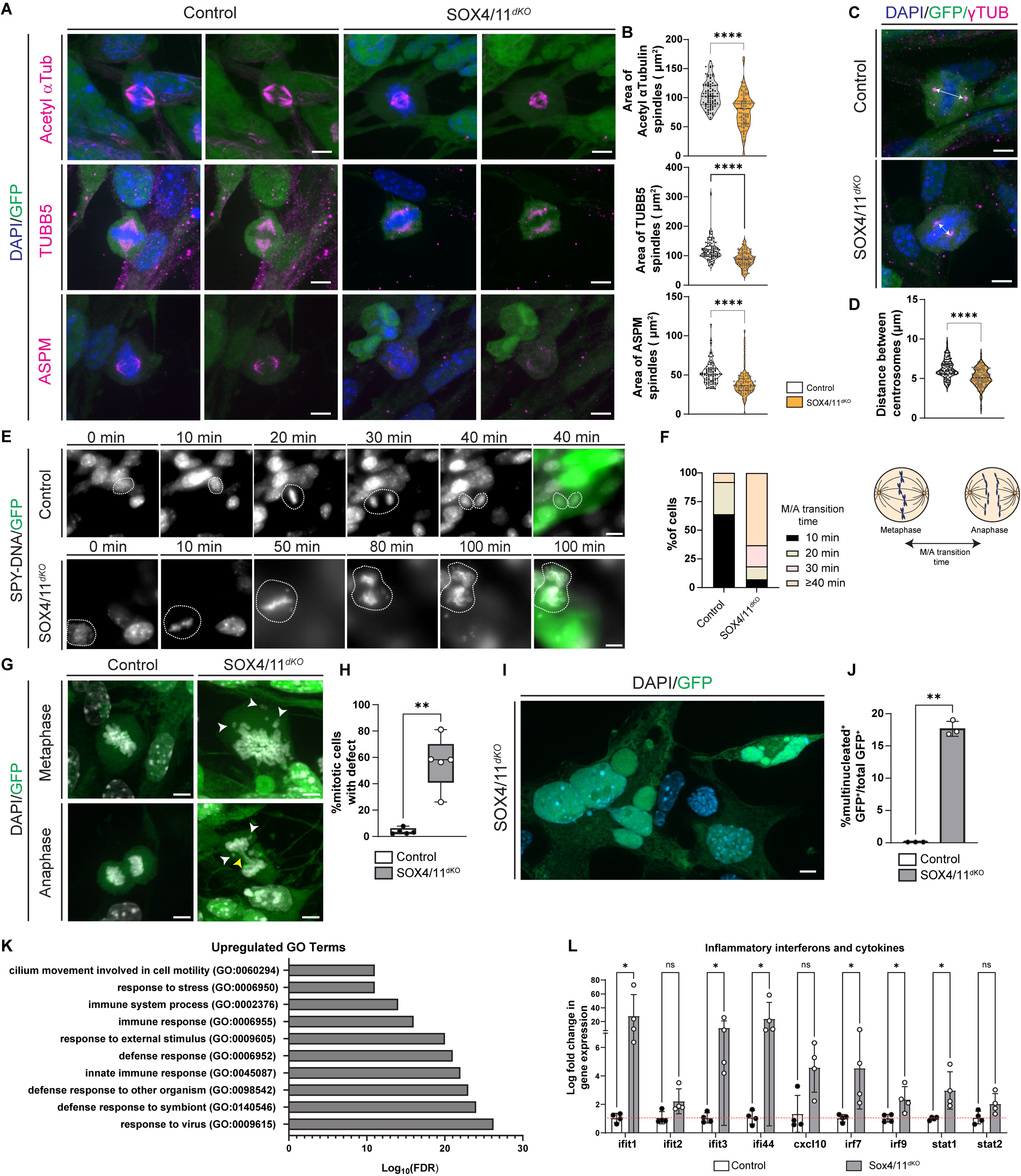
Loss of SOX4/11 from aNSPCs leads to mitotic catastrophe: **(A)** Immunocytochemical analysis of mitotic spindle associated proteins acetyl α-Tubulin, β-Tubulin 5 (TUBB5) and ASPM (magenta) in transduced cells (green). DNA was stained with DAPI (blue) to identify metaphase cells. Scale bar = 5µm **(B)** Quantification of area covered by Acetyl α-Tubulin, β-Tubulin 5 (TUBB5) and ASPM. SOX4/11^dKO^ cells showed a reduced area covered by spindle associated proteins. At least 30 transduced metaphase cells were analyzed per condition. n=3 biological replicates. **(C)** Immunocytochemical analysis of centrosomes using γ-Tubulin staining (γ-Tub, magenta). Spindle length was measured as the distance between the two centrosomes (white arrows) Scale bar = 5µm **(D)** Quantification of spindle length. At least 30 transduced metaphase cells having only 2 centrosomes were analyzed per condition. n=3 biological replicates. **(E)** Frames from live cell imaging of transduced (green) control and SOX4/11^dKO^ cells. SPY-DNA (grey) was used to observe mitotic events. SOX4/11^dKO^ frame was cut from 20 to 50 min for representation purposes, all frames can be found in the supplementary video (Movie EV5.1, Movie EV5.2). Scale bar= 5µm **(F)** Quantification of time taken for cells to transition from metaphase to anaphase. Percentage of cells and their M/A transition time was calculated from the 1st frame where cell appeared in metaphase to 1st frame where the cells were in anaphase as seen in the representation. SOX4/11^dKO^ cells (bottom panel) spent a longer time transitioning from metaphase to anaphase. 10 cells per condition from 3 biological replicates analyzed. **(G)** Immunocytochemical analysis of mitotic transduced cells (green) showing presence of mitotic defects such as the presence of mis-aligned chromosomes (white arrows) in metaphase (upper panel) and mis-segregated chromosomes (white arrows) and DNA bridges (yellow arrows) in anaphase (lower panel) in SOX4/11^dKO^ cells. DAPI in grey. Scale bar = 5µm. **(H)** Quantification of percentage of transduced cells that showed mitotic defects. At least 50 transduced metaphase were analyzed per condition. n=3 biological replicates. **(I)** Representative image showing transduced SOX4/11^dKO^ cells (green) with multiple nuclei (DAPI, in blue). Scale= 5µm. **(J)** Quantification for percentage of transduced cells showing the presence of multinucleated cells. n=3 biological replicates. **(K)** GO term analysis of 109 upregulated genes shows enrichment of terms related to immune response. **(L)** qRT-PCR validation of selected upregulated genes confirms an increase in RNA expression of genes associated with signal transducers and downstream effector interferons. n=4 biological replicates. Data represented as mean ± SD; t test was used to determine significance. ***P<0.001, **P<0.01, *P<0.05, ns = non-significant.

To study the impact of mitotic spindle architecture defects on mitosis dynamics and timing, live-cell imaging of SOX4/11^dKO^ and control aNSPCs labeled with a fluorescent DNA probe was performed. Interestingly, SOX4/11^dKO^ cells lagged behind control cells to complete mitosis and required on average more time to proceed from metaphase to anaphase (Figure 5E, F, Movie EV5.1, 5.2). More than 50% of control cells proceeded from metaphase to anaphase within the first 10 minutes of the observation period and almost 90% of control cells had completed metaphase to anaphase transition within 20 minutes. In contrast, less than 10% of SOX4/11^dKO^ cells proceeded from metaphase to anaphase within the first 10 minutes and more than 60% of SOX4/11^dKO^ cells failed to make the progression from metaphase to anaphase within 40 minutes (Figure 5H), indicating that SOX4/11^dKO^ cells had a lengthened M-Phase.

We next visualized condensed DNA in M-Phase using DAPI staining in order to analyze chromosome alignment in metaphase and segregation during anaphase. Around 60% of mitotic SOX4/11^dKO^ cells showed metaphase defects (misaligned chromosomes) or anaphase defects (lagging chromosomes; DNA bridges), while mitotic defects were virtually absent from mitotic control aNSPCs (Figure 5G, H). Moreover, DNA staining revealed that mitotic defects in SOX4/11^dKO^ cells resulted in the generation of DNA bridges between cells, cells with micronuclei and multinucleated cells (Figure 5I, J).

Mitotic defects can trigger the initiation of a protective mechanism termed mitotic catastrophe that induces cell death to prevent cancerous cell formation or the accumulation of dysfunctional cells (Sazonova *et al*, 2021). In mitotic catastrophe, p53 activation as well as the induction of a cytokine and type I IFN response play central roles. Activation of p53 triggers proliferation arrest to allow for repair of limited DNA damage. In cells with excessive DNA damage or severe mitotic errors, prolonged p53-induced proliferation arrest initiates a cell death program (Contadini *et al*, 2019; Orth *et al*, 2012). Consequences of mitotic errors such as micronuclei and DNA bridges activate a pro-inflammatory cytokine and type I IFN response, which in turn results in recruitment of immune cells and cell death (Flynn *et al*, 2021; Mackenzie *et al*, 2017; Takaki *et al*, 2024). Indeed, we observed multinucleated cells showing p53 immunoreactivity in SOX4/11^dKO^ cells (EV Figure 5C, D), suggesting that SOX4/11 dKO cells harboring mitotic errors activated p53 signaling. Moreover, GO-term analysis of upregulated genes in SOX4/11 dKO cells indicated an enrichment of genes associated with the immune response (Figure 5K). qRT-PCR analysis confirmed that signal transducers (STAT1, STAT2, IRF7, IRF9) and even more prominently effectors of interferon beta signaling such as IFIT1/2/3 and IFI44 had increased expression in SOX4/11^dKO^ cells (Figure 5L). Collectively, these data indicate that SOX4/11-deficiency resulted in mitotic catastrophe induced cell death.

### SOXC deletion is associated with mitotic defects, M-Phase lengthening and microglia activation in the adult hippocampal neurogenic niche

Next, we analyzed proliferating cells in the dentate gyrus of young adult SOX4/11*^iNestin^* mice for evidence of mitotic defects. Staining against pHH3^+^ cells was applied to identify cells in M-Phase. Because of the relatively low number of pHH3^+^ cells in the adult mouse dentate gyrus, only few pHH3^+^ cells in metaphase and anaphase could be analyzed. Nevertheless, misaligned chromosomes, lagging chromosomes and DNA bridges could be observed in SOX4/11*^iNestin^* mice (Figure 6A). The low number of cells in metaphase and anaphase, however, precluded statistically meaningful quantification. Our in vitro data demonstrated that SOX4/11-deficiency linked mitotic defects were paralleled by a prolonged M-Phase or cell-cycle arrest in M-Phase (Figure 5G, H). We analyzed the fraction of proliferating cells in M-Phase (pHH3^+^) amongst proliferating cells to determine, whether SOX4/11-deficiency in vivo also caused prolongation of M-Phase or cell-cycle arrest in M-Phase. To identify cell-cycle defects, we analyzed mice that received BrdU (50mg/kg per day) for 2 days immediately after tamoxifen induction and were sacrificed 3hrs post BrdU injection. Interestingly, the fraction of cells in M-Phase (pHH3^+^) amongst recombined BrdU^+^ cells was significantly increased in SOX4/11*^iNestin^* mice compared to control mice (Figure 6B, C), suggesting M-Phase lengthening/arrest or increased mitotic re-entry of proliferating cells in SOX4/11*^iNestin^* mice. To distinguish between these possibilities pHH3 staining was combined with staining against MCM2, a marker expressed by proliferating cells throughout the cell-cycle. Compared to control mice, the fraction of pHH3^+^ cells amongst MCM2^+^ proliferating cells was significantly higher (Figure 6D, E), indicating M-Phase lengthening/arrest in proliferating SOX4/11-deficient cells in vivo.

**Figure 6:**
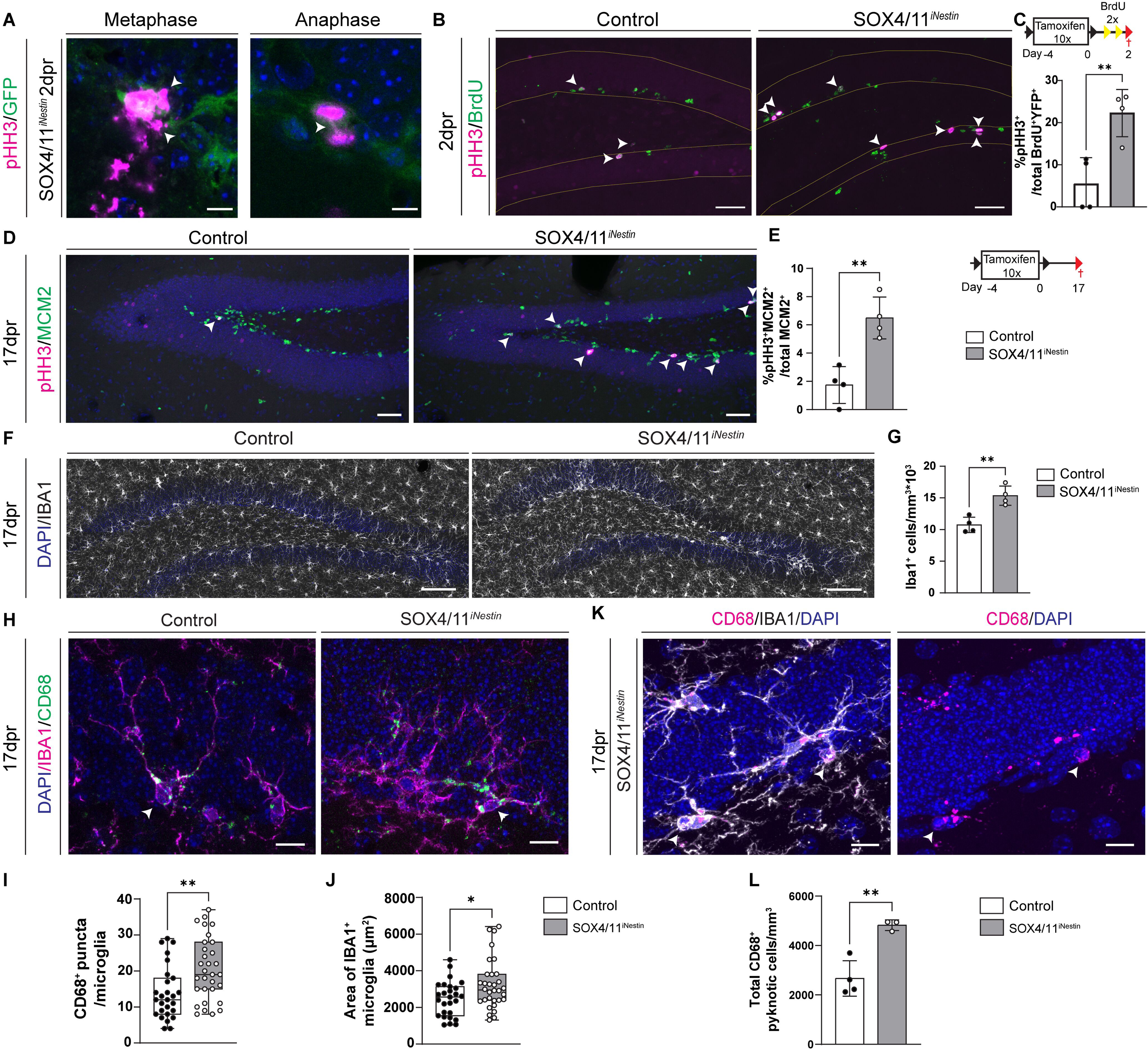
SOX4/11 deletion causes mitotic defects and inflammation in vivo: **(A)** Representative image showing mitotic defects in the dentate gyrus of SOX4/11^iNestin^ mice. pHH3 (magenta) labeling of mitotic DNA revealed misaligned chromosomes in metaphase (left) and DNA bridges (right) in anaphase in recombined cells (green). Scale bar = 5µm **(B)** Immunohistochemical analysis showing an increase in the co-localization (white arrows) of pHH3 (magenta) and BrdU (green) in the dentate gyrus of SOX4/11^iNestin^ mice at 2dpr. Scale bar = 50µm **(C)** Schematic representation of the experimental timeline and quantification of the fraction of BrdU^+^ cells expressing pHH3. n= 4 animals. **(D)** Representative images of the dentate gyrus showing an increase in cells (white arrows) co-expressing pHH3 (magenta) and MCM2 (green) in SOX4/11^iNestin^ mice at 17dpr. Scale bar = 50µm **(E)** Schematic representation of the experimental timeline for 17dpr and quantification of the fraction of MCM2^+^ cells expressing pHH3. n= 4 animals. **(F)** Representative images of the dentate gyrus showing an increase in IBA1^+^ microglia (grey) in SOX4/11^iNestin^ mice at 17dpr. Scale bar= 100µm **(G)** Quantification of IBA1^+^ cells in the subgranular zone at 17dpr. n= 4 animals. **(H)** Representative images showing increased CD68^+^ puncta (green) in IBA1^+^ (magenta) microglia of SOX4/11^iNestin^ mice at 17dpr. DAPI in blue. Scale bar = 10µm **(I)** Quantification of CD68^+^ puncta (≥ 5 µm) in microglia. Minimum of 10 microglia per animal were analyzed. n=3 animals. **(J)** Quantification of IBA1^+^ area of reconstructed microglia. Minimum of 10 microglia per animal were reconstructed and analyzed. n=3 animals. **(K)** Representative image of IBA^+^ microglia (grey) in SOX4/11iNestin mice forming a CD68^+^ pocket (magenta) engulfing pyknotic nuclei. Nuclei are counterstained with DAPI (blue). Scale bar = 10µm. **(L)** Quantification of pyknotic nuclei engulfed by CD68^+^. n=3-4 animals. Data represented as mean ± SD; t test was used to determine significance. ***P<0.001, **P<0.01, *P<0.05, ns = non-significant.

In embryonic neurogenesis, mitotic errors in neural precursor cells induce apoptosis and microglia activation, resulting in rapid clearing of defective cells by phagocytosis (Shi *et al*, 2019). To investigate microglia activation in the hippocampal neurogenic niche, we first quantified the number of microglia in the sub-granular zone (SGZ) of the dentate gyrus using IBA1 as a marker. Indeed, at 17 dpr the number of microglia was significantly increased in SOX4/11*^iNestin^* mice compared to controls (Figure 6F, G). Next, we analyzed the state of activity of microglia using the levels of the microglia-lysosome associated marker CD68 and microglia size as proxies. Notably, IBA1^+^ microglia in the SGZ of SOX4/11*^iNestin^* mice contained significantly more CD68^+^ puncta than microglia in control mice (Figure 6H, I). Moreover, individual IBA1^+^ microglia were larger in SOX4/11*^iNestin^*mice compared to control mice (Figure 6J). Collectively, these data suggest that deletion of SOX4/11 in the neurogenic lineage resulted in a local activation of microglia and an inflammatory response. Importantly, we also observed an increase in the number pyknotic cells that were engulfed by microglial IBA1^+^ processes in SOX4/11*^iNestin^* mice as compared to control mice (Figure 6K, L), suggesting the activation of microglia by dying cells and/or the removal of defective dying cells by microglia in the SGZ.

### SOXC factors are essential for mitotic fidelity in embryonic neurogenesis across species

We next sought to determine whether SOXC factors are also essential for mitotic fidelity in the murine and human embryonic nervous system. In line with previous reports (Thein *et al*., 2010), analysis of published scRNAseq datasets (Delile *et al*, 2019) [Dataref: (Delile *et al*., 2019)] and immunohistochemistry showed SOX4 and SOX11 mRNA and protein, respectively, to be expressed in SOX2 expressing progenitor cells of the E11.5 murine spinal cord (EV Figure 7A-J). We then analyzed Brn4::Cre; SOX4^fl/fl^; SOX11^LacZ/LacZ^ (Brn4^Cre^ SOX4/11^dKO^) transgenic mice (Thein *et al*., 2010), in which SOX4 and SOX11 are deleted from the developing spinal cord. Mice with WT alleles for SOX4 and SOX11 served as controls. In line with previous studies, that had reported substantial cell death of neural precursors in the SOX4/11-deficient mouse neural tube and embryonic spinal cord (Bhattaram *et al*., 2010; Thein *et al*., 2010), we observed numerous pockets of pyknotic nuclei in the E11.5 spinal cord ventricular zone of Brn4^Cre^ SOX4/11^dKO^ transgenic mice (Figure 7A). Analysis of pHH3^+^ mitotic cells in the ventricular zone showed a significant increase in the fraction of cells that harbored mitotic defects such as mis-segregated chromosomes and DNA bridges in Brn4^Cre^ SOX4/11^dKO^ mice compared to controls (Figure 7B, C). In addition, we occasionally observed cells with micronuclei in the spinal cord ventricular zone of Brn4^Cre^ SOX4/11^dKO^ mice (Figure 7D). These observations indicate that loss of SOX4 and SOX11 causes mitotic defects in neural precursor cells of the developing mouse spinal cord.

**Figure 7:**
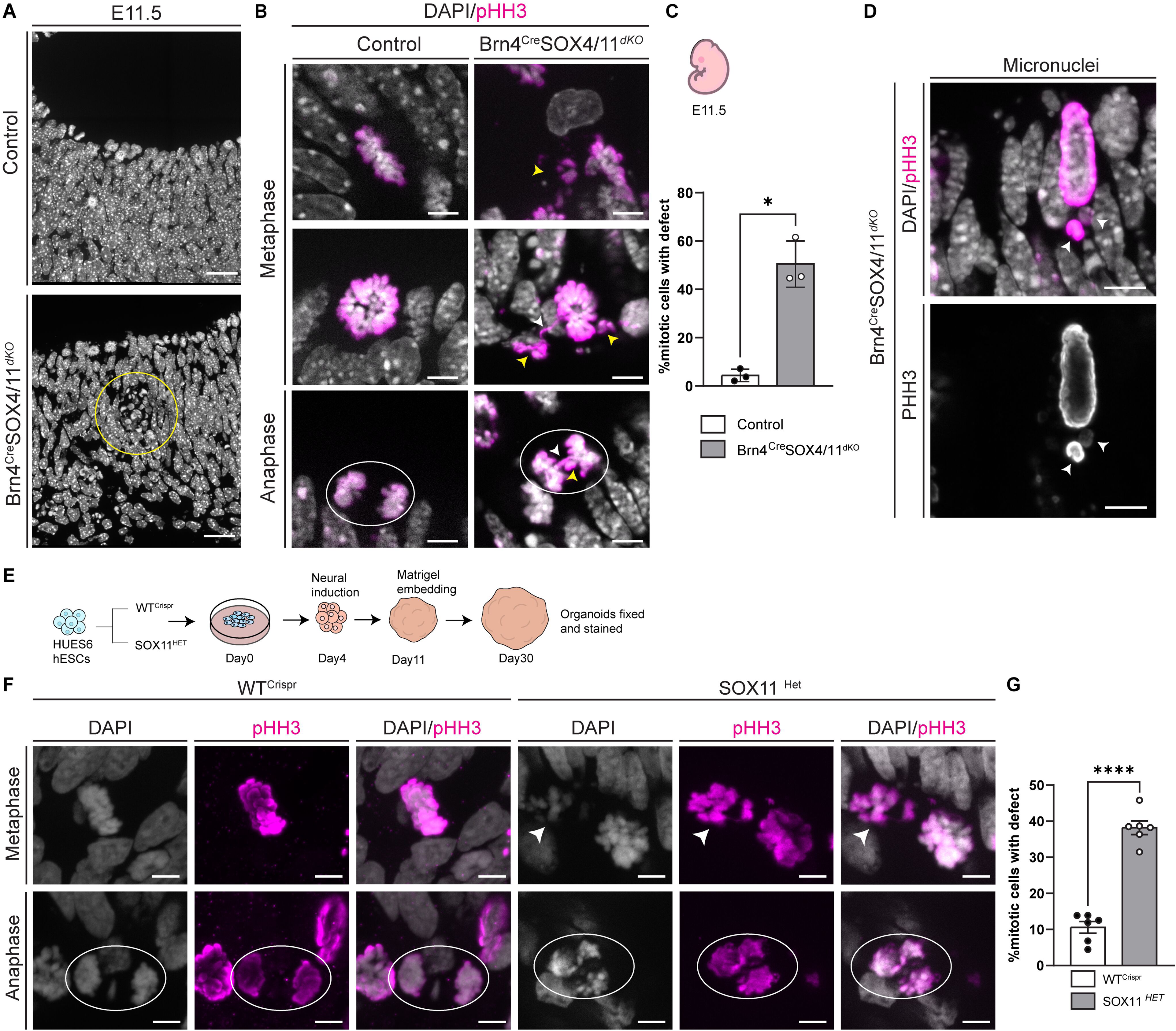
SOXC factors maintain mitotic fidelity in murine and human CNS development: **(A)** Representative images of the E11.5 spinal cord of control and Brn4^Cre^ SOX4/11^dKO^ mice. Note the area of pyknotic nuclei (yellow circle) in close proximity to the ventricle in Brn4^Cre^ SOX4/11^dKO^ mice. Scale bar = 5µm. **(B)** Representative images of the E11.5 spinal cord ventricular zone of control and Brn4^Cre^ SOX4/11^dKO^ mice. pHH3 staining (magenta) was used to identify and analyze mitotic DNA; DAPI labeling (grey) was used as a general DNA label. Note the mitotic defects in E11.5 spinal cord Brn4^Cre^ SOX4/11^dKO^ mice: misaligned chromosomes (white arrows) in metaphase and DNA bridges (yellow arrow) and mis-segregated chromosomes (white arrows) in anaphase. Scale bar = 5µm. **(C)** Quantification of the percentage of mitotic cells exhibiting mitotic defects in control and Brn4^Cre^ SOX4/11^dKO^ spinal cord. A minimum of 50 metaphase or anaphase cells were analyzed per animal. n= 3 animals. Example of a micronucleus in the E11.5 spinal cord ventricular zone of Brn4^Cre^ SOX4/11^dKO^ mice. pHH3 (magenta); DAPI (grey). Scale bar = 5µm. **(E)** Schematic representation of the brain organoid formation protocol used in F and G. **(F)** Representative images of metaphase and anaphase nuclei in the ventricular zone WT^Crispr^ and SOX11^Het^ organoids. Note the chromosome misalignment (white arrows) in metaphase and abnormal anaphase separation (white circle) in SOX11^Het^ organoids. pHH3 staining (magenta) was used to identify and analyze mitotic DNA; DAPI labeling (grey) was used as a general DNA label. Scale bar = 5µm. **(G)** Quantification for percentage of mitotic cells exhibiting mitotic defects in WT^Crispr^ and SOX11^Het^ organoids. A minimum of 40 metaphase or anaphase cells were analyzed. n= 3 organoids from 2 batches. Data represented as mean ± SD; t test was used to determine significance. ***P<0.001, **P<0.01, *P<0.05, ns = non-significant.

In humans, heterozygous loss-of-function mutations in SOX11 have been causally linked to SOX11-syndrome or Coffin-Siris Syndrome 9 - a neurodevelopmental syndrome associated with microcephaly (Al-Jawahiri *et al*, 2022; Hempel *et al*, 2016; Tsurusaki *et al*, 2014). Here, we studied the impact of SOX11 heterozygous knockout on mitotic fidelity of human neural precursor cells. We previously generated SOX11^+/-^ human embryonic stem cells (SOX11^HET^) and isogenic control lines (WT^Crispr^) from the human embryonic stem cell line HUES6 by CRISPR/Cas9 genome engineering (Turan *et al*, 2019). Brain organoids were generated using the protocol by Lancaster and colleagues (Giandomenico *et al*, 2019; Lancaster *et al*, 2013) with slight modifications. Analysis was performed at 30 days post aggregation of embryonic stem cells (Figure 7E). Evaluation of pHH3^+^ mitotic cells in the ventricular zone of neural rosette structures revealed a 4-fold increase in the incidence of mitotic defects in SOX11^Het^ organoids compared to WT^CRISPR^ organoids (Figure 7F, G). Moreover, qRT-PCR analysis showed that SOX11^het^ organoids expressed an inflammatory profile reminiscent of the inflammatory profile observed in SOX4/11-deficient aNSPCs (EV Figure 7K). Collectively, the findings of increased mitotic defects in the SOX4/11-deficient murine embryonic spinal cord and in human SOX11 heterozygote knockout brain organoids indicate that SOXC factors are essential for mitotic fidelity in neural precursor cells across ontogeny and species.

## Discussion

Here, we uncover a surprising novel function for the neuronal fate determining transcription factors SOX4 and SOX11 in maintaining mitotic fidelity of fast proliferating precursor cells in the adult hippocampus. Deletion of SOX4/11 caused severe mitotic defects in neural precursor cells as evidenced by prolongation of M-Phase, mitotic spindle and centrosome defects, misaligned chromosomes, lagging chromosomes and DNA bridges, as well as the generation of progeny with micronuclei and multinucleation. Our data identify mitotic catastrophe – a mitosis defect-induced cell death mechanism that prevents the accumulation of cells with genomic defects (Sazonova *et al*., 2021) as the underlying cause for death of SOX4/11-deficient precursor cells. In line with the activation of mitotic catastrophe, we find increased activity of the p53 pathway and expression of a cytokine and type I IFN response in SOX4/11-deficient precursor cells.

Previous findings indicate that mitotic errors in neural precursor cells of the developing nervous system not only trigger mitotic catastrophe but also microglia activation, resulting in rapid clearing of defective cells by phagocytosis (Shi *et al*., 2019). Consistent with the rapid clearing of mitotically defective IPCs, we found only few cells with mitotic defects but enhanced microglial activation and increased engulfment of recombined and pyknotic cells in the dentate gyrus of adult SOX4/11*^iNestin^* mice. In the E11.5 mouse spinal cord, where fast dividing precursor cells are far more abundant than in the adult mouse dentate gyrus and where microglia have only begun to invade the nervous system parenchyma (Rigato *et al*, 2011), dividing cells with mitotic defects were frequently observed following deletion of SOX4/11. This observation further supports the idea that SOX4/11 control mitotic fidelity of neural precursor cells in vivo and that microglial phagocytosis rapidly removes precursor cells with mitotic defects.

Adult SOX4/11*^iNestin^* mice showed increased activation of RGLs and generation of IPCs. This observation may suggest that SOX4 and/or SOX11 participate in the regulation of RGL quiescence and activation in the adult dentate gyrus. Enhanced activation of RGLs and generation of IPCs, however, was observed in both the recombined and the non-recombined RGL-lineage. This observation argues that increased RGL activity may in fact reflect an indirect effect of SOX4/11 deletion rather than a cell-autonomous regulatory function of SOX4/11 in RGL quiescence and activation. Indeed, it has been suggested that immature neurons provide negative feedback signals onto RGLs to restrict their activity (Hodge *et al*, 2012). Considering that SOX4/11-ablation resulted in a substantial reduction in newborn neurons, increased activation of RGLs and generation of IPCs may be the consequence of decreased negative feedback from immature neurons onto RGLs.

The present study cannot distinguish the individual contribution of SOX4 and SOX11 to the mitotic spindle phenotype. Some studies demonstrated that SOX4 and SOX11 operate redundantly as critical regulators of precursor cell survival in embryonic mouse development (Bhattaram *et al*., 2010; Thein *et al*., 2010); while other findings suggested that SOX4 and SOX11 fulfill non-redundant functions in the regulation of cortical neural precursor cell proliferation and maintenance (Chen *et al*., 2015; Wang *et al*., 2013). Future studies using single SOX4 and SOX11 knockouts should directly address the function of these proteins in safeguarding mitotic fidelity.

It is interesting to note, that in human ESC-derived brain organoids loss of a single functional SOX11 allele is sufficient to increase the frequency of mitotic defects in the ventricular zone. This observation indicates that SOX11 is critical for safeguarding mitotic fidelity in human neural precursor cells and suggests that loss of SOX11 may be responsible for the mitotic defects in SOX4/SOX11-deficient mouse neural precursor cells. SOX11-haploinsufficiency has been repeatedly found to be associated with microcephaly (Al-Jawahiri *et al*., 2022; Hempel *et al*., 2016; Schincariol-Manhe *et al*, 2024), a neurodevelopmental phenotype frequently observed in pathogenic variants of genes linked to mitotic spindle function (Phan & Holland, 2021). The present data raise the interesting idea that SOX11-linked microcephaly is caused by impaired function of the mitotic spindle in neural precursor cells.

How SOXC factors control the function of the mitotic spindle remains to be determined. Previous studies described SOXC factors as important survival factors for neural and mesenchymal precursors in mouse embryogenesis. Moreover, the study by Bhattaram and colleagues (Bhattaram *et al*., 2010) provided evidence that in the case of mesenchymal precursors SOXC factors control cell survival at least in part through the regulation of Tead2 and the Hippo pathway. In that study, transcriptomic analysis of SOX4/11-deficient tissue enriched for mesenchymal progenitor cells, however, did not find dysregulation of mitosis-related genes, suggesting that cell death of SOX4/11-deficient mesenchymal precursor cells and cell death of SOX4/11-deficient neural precursor cells may be caused by different mechanisms. Our data indicate that in adult neural precursor cells SOXC factors control the expression of genes encoding for essential proteins in microtubule dynamics, the centrosome, the spindle-assembly checkpoint and cytokinesis. In general, expression of G2/M phase and mitotic genes is controlled by a core transcription factor complex consisting of the transcription factors B-MYB, FOXM1 and the MuVB complex (Fischer *et al*, 2022). Whether SOXC factors interact with this transcription factor complex to control mitotic fidelity remains to be determined. Mitosis is a fundamental cell biological process in all dividing cells; SOXC expression, however, is restricted to select populations of precursor cells including neural precursor cells. It will be interesting to investigate why neural precursor cells depend on SOXC transcription factors for proper execution of mitosis. Recent studies discovered that the centrosome as a key organizing center for cytoskeleton and the mitotic spindle apparatus features substantial differences in its proteome composition between cell-types (Pilaz *et al*, 2016). It is tempting to speculate that SOXC proteins as neurogenic transcription factors contribute to setting up a neural precursor-specific mitotic spindle apparatus. Wiring transcriptional neuronal fate determinants such as SOXC proteins into the mitosis machinery may allow to precisely time different phases of the cell-cycle and the duration of mitosis, which have been shown to have a major impact on cell fate choices of neural precursors (Arai *et al*, 2011; Lange *et al*, 2009; Mitchell-Dick *et al*, 2019; Pilaz *et al*., 2016).

In adult hippocampal neurogenesis about three out of four fast proliferating IPCs undergo cell death and are removed by microglial phagocytosis (Kempermann *et al*, 2003; Pilz *et al*., 2018; Sierra *et al*, 2010). The cause for this substantial amount of cell death amongst IPCs remains unknown, but it has been suggested that IPC mitosis and cell death are functionally related (Sierra *et al*., 2010). A recent study reported that SOXC factor stability in mitotic embryonic neural precursor cells is actively controlled by GSK3β-dependent phosphorylation (Da Silva *et al*, 2021). It will be interesting to investigate, whether differences in the SOXC phosphorylation/stability pathway and SOXC-dependent mitotic fidelity drive the decision between IPC survival and cell death in the adult hippocampal neurogenic niche.

## Material and methods

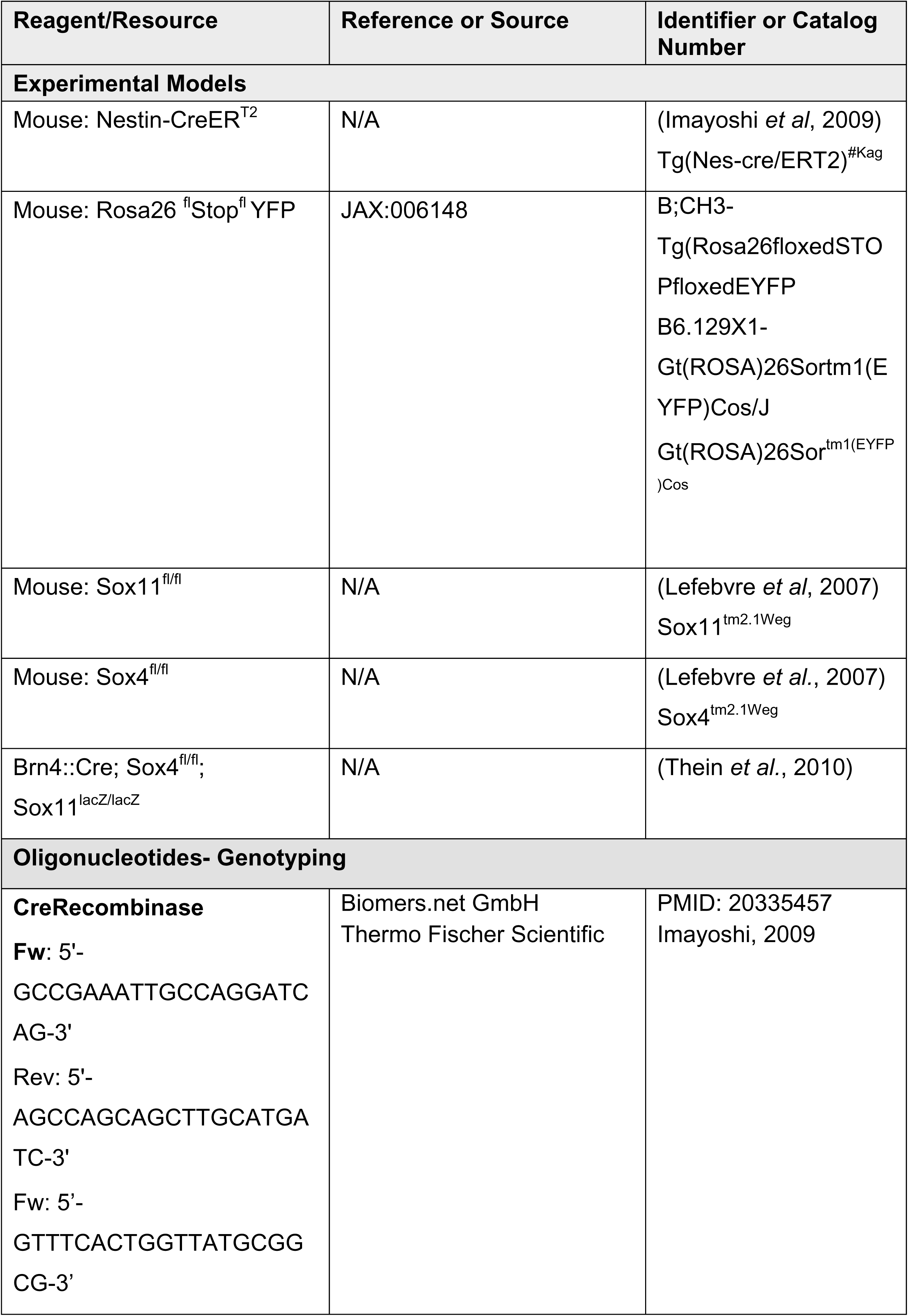
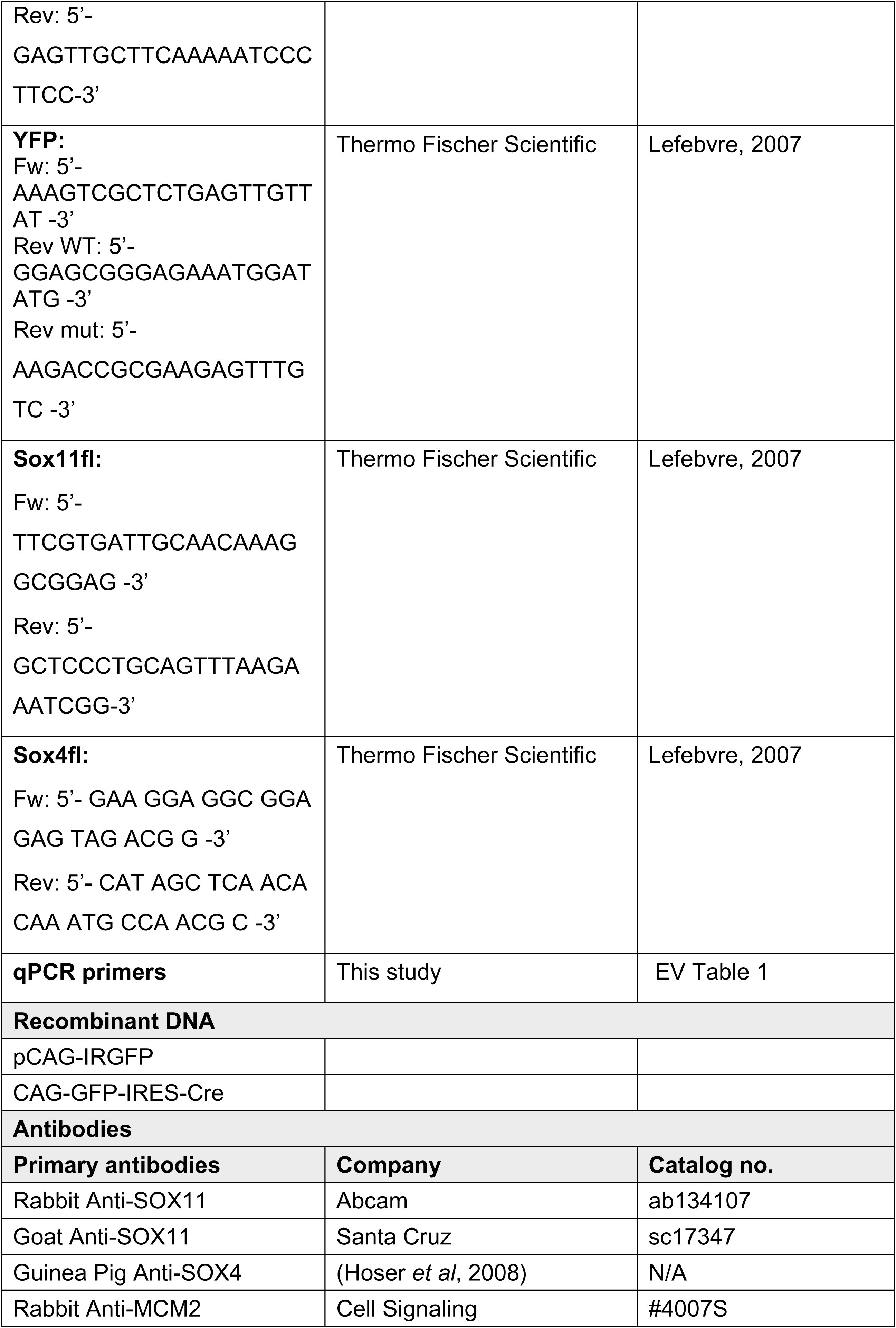
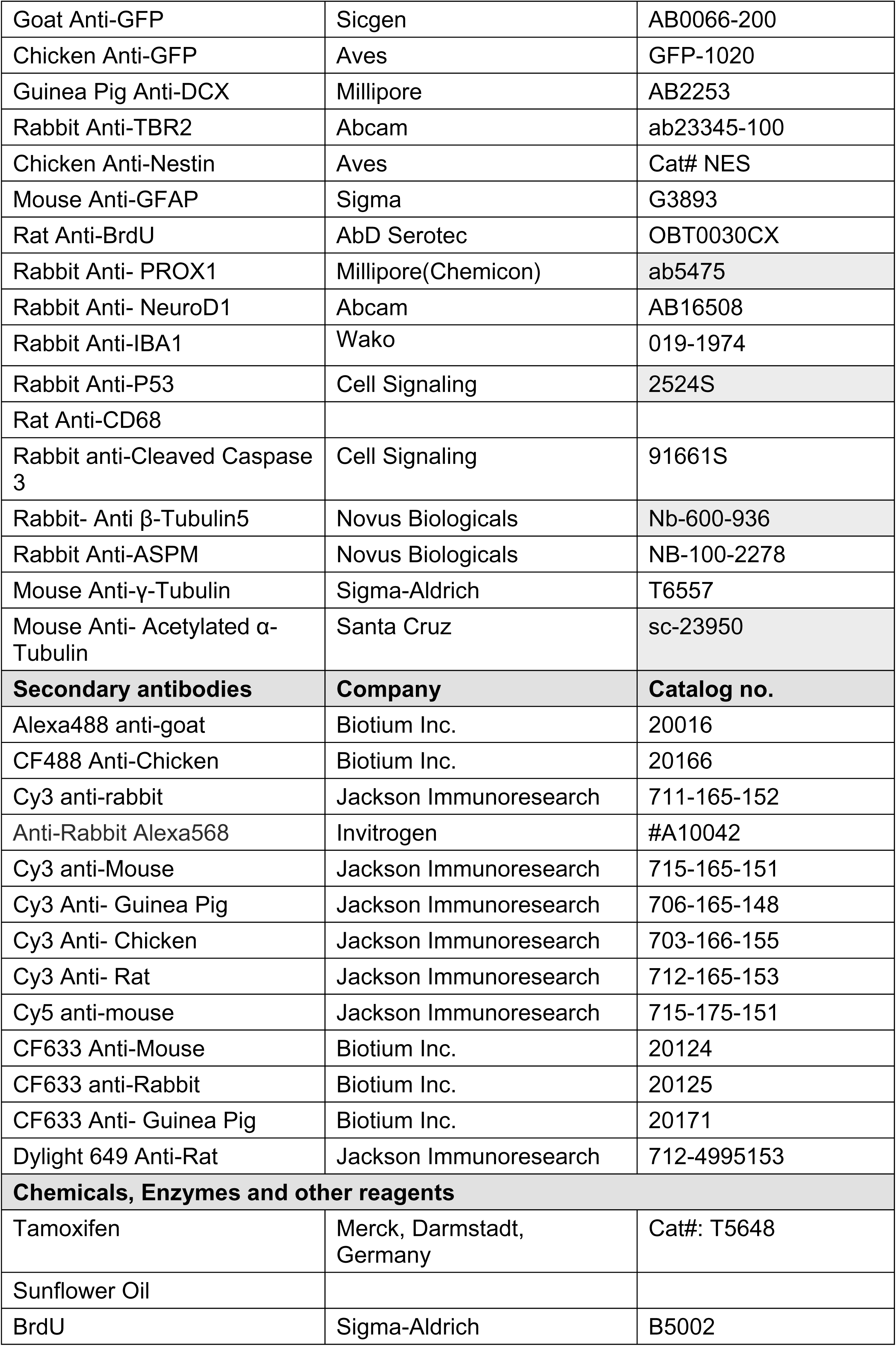
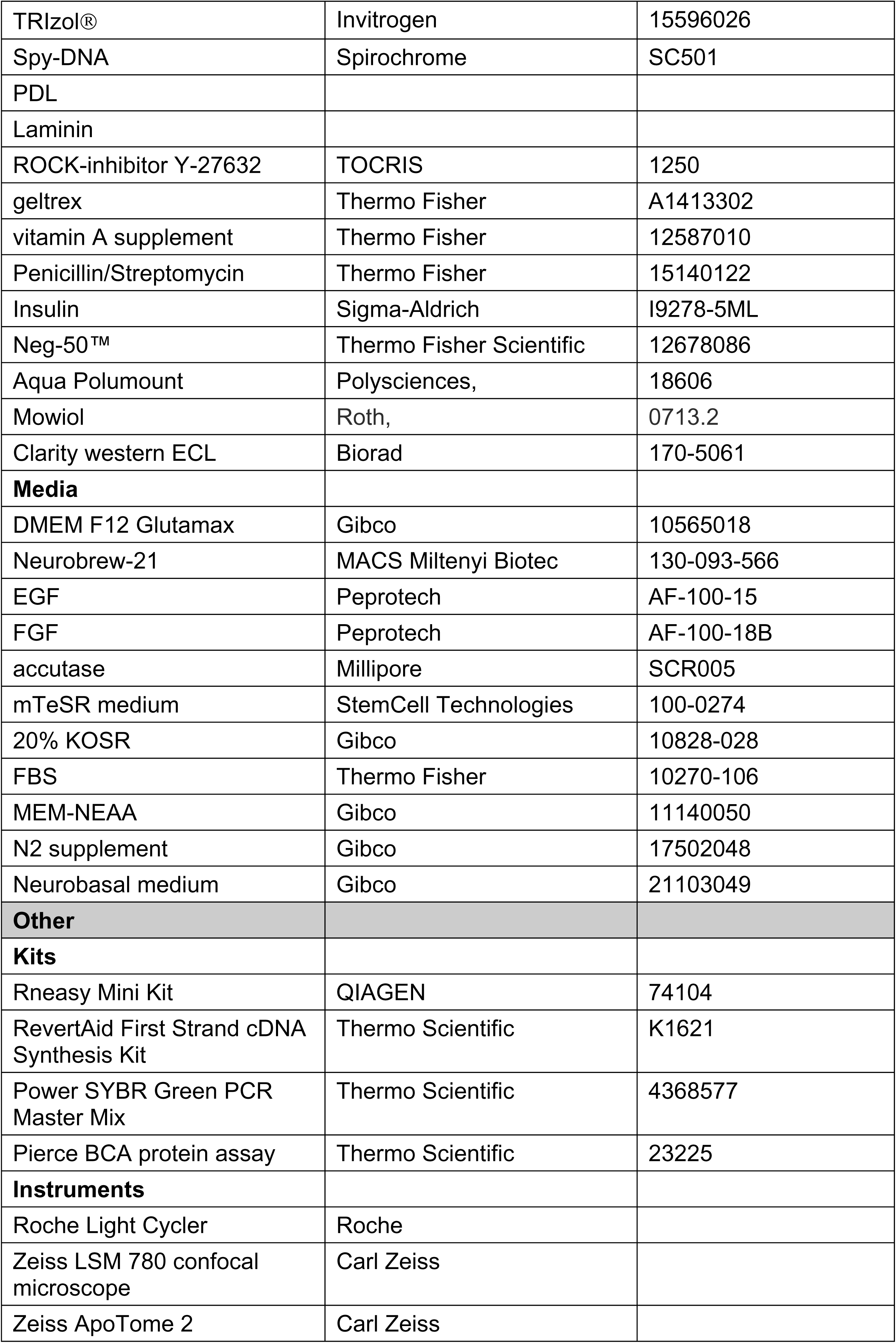
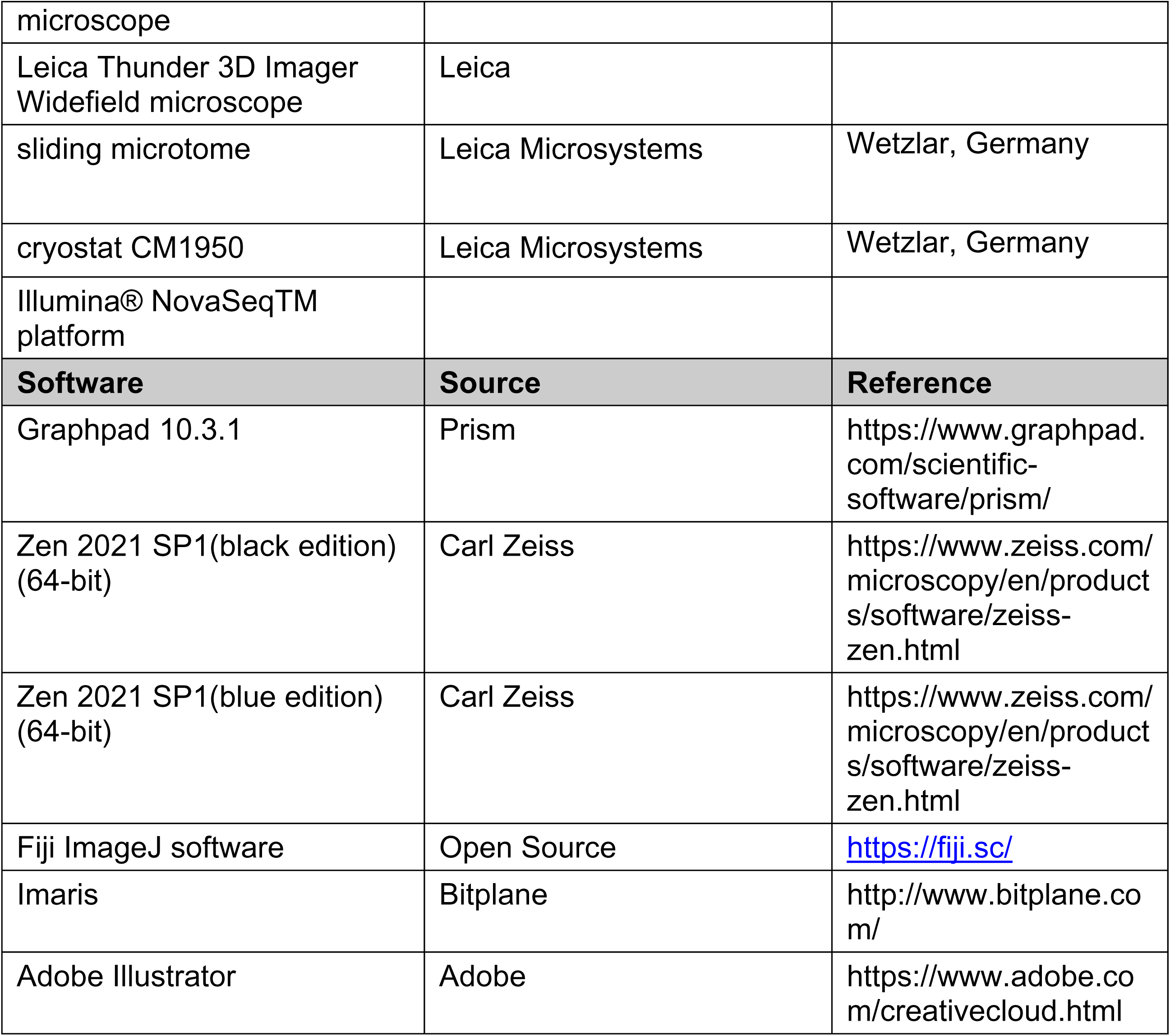

### Mouse model

All experiments were carried out in accordance with the European Communities Council Directive (86/609/EEC) and were approved by the government of Middle-Franconia. Nestin::CreER^T2^ (Imayoshi *et al*., 2009), Rosa26floxedSTOPfloxedEYFP (B;CH3-Tg(Rosa26floxedSTOPfloxedEYFP; JAX:006148) Sox4^flox/flox^ [(Lefebvre *et al*., 2007) Sox4^tm2.1Weg^ and Sox11^flox/flox^ [(Lefebvre *et al*., 2007) Sox11^tm2.1Weg^] have been described previously. Nestin::CreERT2; Rosa26floxSTOPfloxYFP; Sox4flox/flox Sox11flox/flox animals were initially generated from the same cross, and subsequently maintained as separate lines in the following to enable homozygous breeding. Male and female mice were used for experiments. For all experiments, mice were grouped housed in standard cages with ad libitum access to food and water under a 12h light/dark cycle.

For embryonic spinal cord analysis, Brn4::Cre; Sox4fl/fl; Sox11lacZ/lacZ mice previously described (Thein *et al*., 2010) were used. Mice with WT alleles for SOX4 and SOX11 served as controls.

### Genotyping

Genotyping PCR was performed using the aforementioned primers for Cre, YFP and SOX4 and SOX11 (see primer table).

### Tamoxifen injections

Tamoxifen was dissolved in ethanol and sunflower seed oil (1:9) to a concentration of 10 mg/ml. For induction of recombination animals were intraperitoneally (i.p.) injected with 34mg/kg bodyweight Tamoxifen every twelve hours for five consecutive days (Mori *et al*, 2006).

### BrdU injections

Bromodeoxyuridine (BrdU) was dissolved in 0.9% NaCl at a concentration of 10 mg/ml. Animals were intraperitoneally injected with 50 mg/kg bodyweight BrdU either once a day for two consecutive days following the tamoxifen injections or five times a day at 3hr intervals on day 17 after the last tamoxifen injection.

### Tissue processing

Mice were killed using CO2 and were transcardially perfused with phosphate-buffered saline (pH7.4) for five minutes at a flowrate of 20 ml/min. Fixation was carried out by perfusion with 4% paraformaldehyde (PFA) dissolved in 0.1 mM Phosphate Buffer (pH 7.4) for five minutes at a flowrate of 20 ml/min. Brain were removed and stored overnight at 4°C in 4% PFA for post-fixation, followed by dehydration in 30%sucrose in 0.1M phosphate buffer. Frozen brains were coronally sectioned into 40µm thick sections using a sliding microtome (Leica Microsystems, Wetzlar, Germany).

Embryonic spinal cord tissue was collected at day E11.5. in ice cold 1x Phosphate-buffered saline (1x PBS – 0.1 M Na_2_HPO_4_, 0.1 M NaH_2_PO_4_, 0.15 M NaCl) (pH 7.4). Tissue samples were taken for genotyping and embryos were fixed for 2.5 hours in 4 % PFA dissolved in PBS at 4 °C while mechanically shaken. Embryos were washed in 1x PBS followed by over-night incubation in 15% Sucrose (in 0.12 M phosphaste buffer) for cryoprotection. After embryos had sunk in the sucrose solution, the tissue was deep-frozen in gelatin solution (15% Sucrose, 7.5% gelatin dissoloved in 0.12M PB) and stored at -80 °C. 12 – 14 µm sections were cut on Leica cryostat CM1950. Tissue sections were subsequently stored at -80 °C.

### Human brain organoid formation and processing

Media compositions:

<u>OFM</u>: DMEM/F12 + GlutaMAX-1x, 20% KOSR, 3% FBS, 0.1 mM MEM-NEAA, 0.1 mM 2-mercaptoethanol

<u>NIM</u>: DMEM/F12 + GlutaMAX-I, 1% N2 supplement 0.1mM MEM-NEAA, and 1µg/ml Heparin

<u>ODM</u>: 1:1 mix of DMEM/F12 + GlutaMAX-I and Neurobasal medium, 0.5% N2 supplement, 0.1 mM MEM-NEAA, 100 U/ml penicillin and 100 µg/ml streptomycin, 1% B27 +/-vitamin A supplement, 0.025% insulin, 0.035% 2-mercaptoethanol

The generation of SOX11 heterozygote knockout embryonic stem cells and of control embryonic stem cell lines have been previously described (Turan *et al*., 2019) Cells were maintained in mTeSR medium on geltrex (4.8 mg/ml in DMEM/F12+Glutamax;) coated 6-well plates at 37°C until the cells were 80-90% confluent. For generation of brain organoids the Lancaster protocol (Giandomenico *et al*., 2019; Lancaster *et al*., 2013) was used with some modifications. In short, hESCs were split upon reaching confluency using accutase (Millipore, Cat# SCR005) to obtain single cell suspension. Cells were centrifuged at 300 g for 3 minutes and resuspended at a dilution of 8x10E4 cells / ml in organoid formation medium (OFM) with 50 µM ROCK-inhibitor Y-27632 and 4ng/ml of bFGF . 1.2x10E4 cells / well were seeded into 96-well ultra-low attachment plates (Corning #7007) for formation of embryoid bodies. After 72h in culture, fresh OFM lacking FGF and ROCKi was used to replace half the medium. OFM was replaced with neural induction medium (NIM) after 5 days to induce neural differentiation. Media was replaced every 48 h. After 11days in culture the neural aggregates were embedded in matrigel droplets and transferred to organoid differentiation medium (ODM) without vitamin A and incubated in a 10cm petri-plate (Greiner bio-one, # 664-160). After 3 additional days in culture, medium was changed to organoid differentiation media (ODM) containing vitamin A. Plates were kept on an orbital shaker (30rpm). Medium was changed twice per week. Organoids were collected for fixation on day 30.

For fixing the organoids, organoids were collected in a 1.5 ml tube, washed once with 1xPBS, and fixed in 4% PFA in 1xPBS for 30mins. Organoids were washed thrice in 1xPBS and transferred to 30% sucrose in PBS and stored at 4°C. Organoids were embedded into Neg-50™ Frozen Section Medium, frozen on dry ice, and stored at - 80°C. 20µm thick cryosections were made using a Leica cryostat CM1950 and stored at -20°C.

Organoids from the control and heterozygous lines that were generated at the same time were considered as one batch. Analysis was done from two independent batches with 3 organoids from each batch.

### Immunofluorescent staining

Free floating sections were washed thrice for 10mins each in PBS and blocked for 1hr at room temperature in PBS containing 3% donkey serum and 0.25% Triton-X (PBS++). Sections were incubated with the primary antibody diluted in PBS++ for 72h at 4°C, followed by three washes in PBS and incubation with secondary antibodies diluted 1:1000 in PBS++ overnight at 4°C. Sections were washed thrice in PBS followed by a wash in 4′,6-Diamidin-2-phenylindol (DAPI) diluted in PBS (1:10,000) and washed thrice in PBS for 10mins. Sections were mounted in aqua-Poly/Mount ().

For BrdU immunostainings, sections were washed thrice for 10 mins each in PBS, incubated in 2N HCl at 37°C for 10mins, followed by two 10 minute washes with 0.1M Borate and three washes with PBS before proceeding with the staining protocol.

For staining of embryonic spinal cord sections, slides were warmed to room temperature for 5-10 min, washed three times with 1x PBS at 42°C for 5 minutes each, and incubated in blocking solution (0.1% Triton X-100 in 1x PBS (0.1% PBS-T) and 1% BSA) (PBS-T++) for 30 minutes to 1 hour. After blocking, section were incubated with primary antibody diluted in PBS-T++ overnight at 4°C. The next day, samples were washed thrice for 30 minutes each in PBS-T and incubated with secondary antibodies diluted in PBS-T++ for 1 hour at room temperature. Samples were washed three more times in PBS-T and were counterstained with 4′,6-diamidine-2′-phenylindole (DAPI). Sections were mounted in Mowiol solution (10% w/v Mowiol, in 0.1M TRIS-HCl, pH 8.5 and 25% (w/v) glycerol)).

For organoid sections the slides were brought to room temp and washed thrice with 1xPBS for 5min each and incubated in blocking solution (10%NGS, 3%BSA in 1xPBS+0.1%Tween) for at least 1hour at room temperature. Following the blocking step the slides were incubated in primary antibody diluted in blocking solution, overnight at 4°C. Next day, the slides were washed thrice in 1xPBS for 5 min each and incubated in secondary antibody diluted in blocking solution and incubated for 2h at room temperature. The sections were then counterstained with DAPI and washed twice in 1xPBS before mounting with Mowiol solution (10% w/v Mowiol, in 0.1M TRIS-HCl, pH 8.5 and 25% (w/v) glycerol).

### Adult neural stem/precursor cell (aNSPC) cultures

Adult neural stem and progenitor cells (aNSPCs) were isolated from the dentate gyrus of 8-9 week old SOX4/11^fl/fl^ mice using previously published protocol (Schaffner *et al*, 2018). Cells were cultured as neurospheres in DMEM F12 Glutamax medium supplemented with 1xNeurobrew-21, 8mM HEPES, 10ng/mL EGF and 10ng/mL FGF. The cultures were maintained as free-floating neurospheres in culture flasks (Greiner Bio-one, Cat# 658175).

### Retrovirus production and transduction

The CAG-GFP and CAG-GFP-IRES-Cre MML-retroviruses (Steib *et al*, 2014; Tashiro *et al*, 2006) were produced as previously described (Tashiro *et al*., 2006). For virus transduction, the neurospheres were dissociated into single cells using accutase. For immunofluorescence, 1x10^5^ cells were seeded on PDL/Laminin (50µg/ml PDL and 5µg/ml laminin, coated according to manufacturer’s protocol) coated coverslips in 24 well plates (Greiner bio-one, 662-160) to form a monolayer. For RNA and Protein extraction 1x10^6^ cells were seeded on PDL/Laminin coated 6-well plates (Greiner bio-one, 657-160). Twenty-four hours after seeding, cells were transduced with retrovirus at MOI of 50. Medium was exchanged 24h after transduction. Cells were analyzed 72h after transduction.

### Immunocytochemistry

Coverslips were fixed in 4% PFA for 10mins at room temperature followed by three washes with PBS. The coverslips were incubated for 30mins in PBS++ at room temperature, followed by incubation with the primary antibody diluted in PBS++ overnight. Coverslips were then washed three times with PBS, incubated with secondary antibodies diluted 1:1000 in PBS++ for 2h at room temperature. Coverslips were incubated in DAPI (1:10,000 in PBS) for 10mins and mounted using aqua-Poly/Mount. Coverslips were stored at -20°C.

### RNA isolation and qRT-PCR

RNA isolation was performed using a protocol combining TRIzol® Reagent and the Rneasy Mini Kit (Schaffner *et al*, 2023). For cDNA preparation the RevertAid First Strand cDNA Synthesis Kit was used according to manufacturer’s protocol. qRT-PCR was conducted on a Roche Light Cycler (384-well format) using Power SYBR Green PCR Master Mix according to the manufacturer’s instructions. For each experimental group three independent biological replicates were analyzed in triplicates. For quantification, the 2^-ΔΔCT^ method was applied. Mean CT values of each target gene were normalized to the mean of the mean CT values of two housekeeping genes (Hprt, RPL27). Fold changes were calculated by normalizing each ΔΔCT value to the control ΔΔCT values. Primer sequences are listed in table S1. Primers for qPCR were either self-designed or taken from primer bank (https://pga.mgh.harvard.edu/primerbank/).

### Bulk RNA Sequencing

For RNA sequencing 1x10^6^ cells were seeded on PDL/Laminin coated 6 well plates. 3 independent biological replicates were transduced with CAG-GFP and CAG-GFP-IRES-Cre virus 24h after seeding. RNA was extracted after 72 h after virus transduction. From each sample 2µg of RNA was sent to GENEWIZ Germany GmbH for bulk RNA-seq using the Illumina® NovaSeqTM platform (2 × 150 bp configuration, 20 M paired-end reads per sample). Sequencing data was processed as previously published (Schaffner *et al*., 2023) using the galaxy server. Differentially expressed genes (DEGs) were filtered using an adjusted p-value of 0.05 and log_2_-fold change of >1. As parameters for identification of differentially expressed genes an Benjamini–Hochberger adjusted *p*-value of 0.05 and FDR of 0.05 were applied. Gene Ontology (GO) term analysis was performed using Panther (https://geneontology.org/). RNA-seq data can be accessed in the NCBI gene expression and hybridization array data repository (GEO) database and have the accession number GSE309722.

### SDS PAGE and Western-blot

For protein extraction, cells were washed once in PBS, resuspended in RIPA buffer (50 mM Tris-HCl, pH 8.0, 150 mM NaCl, 1% Nonidet P-40, 0.5% Na-deoxycholate [Fluka BioChemika, cat#30970] 0.1% SDS, 2 mM EDTA, complemented with protease- and phosphatase-inhibitors) and lysed on ice for 20mins. Cells were centrifuged at 2000g for 10 mins at 4°C and the supernatant was collected. The total protein concentration was measured using a Pierce BCA protein assay (Thermo Scientific, Cat# 23225). 30µg of protein were separated on 4-12% Bis-Tris gels (Life Technologies, Cat# NP0322BOX) and transferred to a nitrocellulose membrane. The membrane was blocked using 1% BSA in PBS-0.1%Tween20 (PBST) solution. Membranes were incubated at 4°C overnight with primary antibodies diluted in 5%BSA-PBST. Following 3 washes in PBST, membranes were incubated for 2h at room temperature with the secondary antibody diluted in 5% BSA-PBST. Chemiluniminiscence was visualized using Clarity Western ECL substrate and quantified using the FusionFX (Peqlab, Erlangen, Germany) software. All proteins amounts were calculated relative to β-Actin expression.

### Live cell imaging

For live cell imaging 20,000 cells were seeded in 8 chamber (2.2cm^2^ wells) glass slides (Ibidi Cat#80806) or 40,000 cells µ-Dish ^35mm, high^ Glass bottom petri plates (Ibidi cat no# 81151) coated with PDL/laminin. Cells were transduced with retrovirus 24h after seeding. Medium was exchanged the next day. In order to increase the number of cells in mitosis during the imaging period, cells were synchronized 48 h after virus transduction by incubation with 0.5mM thymidine solution in DMEM++. 24h later cells were released from thymidine block, by washing three times with DMEM/F12 Glutamax. Cells were subsequently cultured in DMEM++. 5h after thymidine release, cells were incubated with Spy-DNA (1:10,000,) diluted in DMEM++. 30mins later cells were imaged using a Leica Thunder 3D Imager Widefield microscope equipped with two lasers (488 and 633nm) and a 63x objective (OICE, FAU-Erlangen-Nürnberg). Live cell imaging was performed for at 10min time intervals with Z step size of 0.5 µm for a stack of 10µm. The images were analyzed frame by frame using ImageJ software to determine the time taken for metaphase to anaphase transition. For each experimental group, a minimum of 10 cells / replicate from 3 biological replicates were analyzed.

### Imaging and analysis

For analysis of hippocampal tissue, Z-stacks were taken with a Zeiss LSM 780 confocal microscope or a Zeiss ApoTome 2 microscope using a 20x objective or 63x oil objective. Expression of stage-specific markers was analyzed using the Zen black software in two sections / animal (n=3-4 per experimental group).

In-Vitro images were taken using Zeiss ApoTome 2 microscope with an 63x objective. 3D reconstruction and area analysis was carried out using the Imaris software (Bitplane AG, Zürich, Switzerland). A minimum of 30 cells / replicate from at least 3 independent biological replicates per experimental condition were used to analyse mitosis, spindle and centrosomal defects.

For analysis of the mouse embryonic spinal cord (n=3/group), Z-stacks were taken on a Zeiss LSM 780 confocal microscope using a 63x objective. In each sample, > 40 cells in metaphase or anaphase in the ventricular zone were analyzed.

For analysis of human brain organoids, Z-stacks were taken on a Zeiss LSM 780 confocal microscope using a 63x objective. In each sample, >40cells in metaphase or anaphase in the ventricular zone were analyzed. Per genotype, 3 organoids from 2 independent batches were analyzed.

### Statistical Analysis

Statistical analysis for all the data was performed using GraphPad (version 10.5.0). Normal distribution of data was determined using the Shapiro-Wilks test. Significance was analyzed either using two-tailed Mann-Whitney U test or Welsh’s t-test depending on the distribution of the data. Differences were considered statistically significant at *p<0.05, **p<0.01 and ***p<0.001. All data are presented as mean ± SD (standard deviation).

## Acknowledgments

We thank Drs. Ruth Beckervordersandforth, Sven Falk, Sebastian Jessberger, Marisa Karow, Tomohisa Toda and all members of the Lie laboratory for helpful discussions. This work was supported by grants from the German Research Foundation (LI 858/9-2 and LI 858/11-1 to D.C.L.; project number 455354162 to A.S.) and the Interdisciplinary Centre for Clinical Research (IZKF) of the University Clinic Erlangen (IZKF Synergy grant to D.C.L. and A.S.). S.A. and A.L.L are members of the research training group 2162 ‘Neurodevelopment and Vulnerability of the Central Nervous System’ of the Deutsche Forschungsgemeinschaft (DFG GRK2162/2). The authors thank the Optical Imaging Centre Erlangen of the FAU for technical and methodological support.

## Author contributions

Conceptualization, S.A., D.C.L.; Investigation, S.A., B.M.H., I.S., A.L.L., S.T., S.K.S., A.S.; Formal analysis, S.A., A.L.L., S.K.S., A.S., D.C.L.; Resources and Funding acquisition, A.S., E.S., D.C.L.; Writing-Original draft and Editing, S.A., D.C.L.

## Data Availability

RNA-seq data can be accessed in the NCBI gene expression and hybridization array data repository (GEO) database and have the accession number GSE309722.

---

**Table EV1: Primer sequences for qRT-PCR.**

**Figure EV 2:**
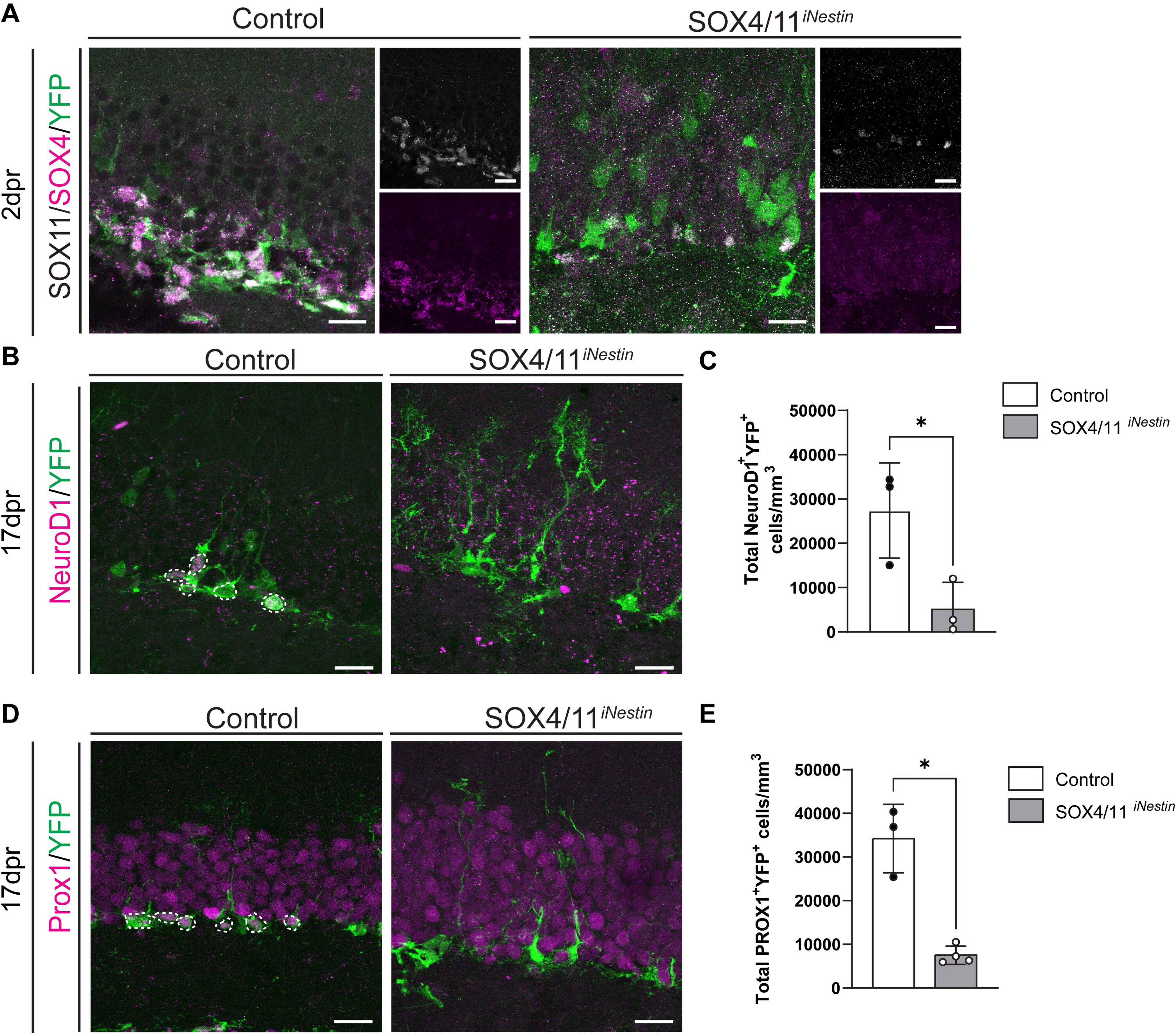
Data supporting Fig 2: **(A)** Representative images of control and SOX4/11^iNestin^ mice at 2dpr. Recombined cells were identified by YFP (green) expression. SOX4 (magenta) and SOX11 (grey) immunoreactivity is lost in YFP^+^ cells in SOX4/11^iNestin^ mice. Scale bar = 20µm. **(B)** Representative images showing loss of NeuroD1 (magenta) expression in recombined cells (green) in SOX4/11iNestin mice 17dpr. Scale bar= 20µm **(C)** Quantification of NeuroD1^+^ YFP^+^ cells in the sub-granular zone of the dentate gyrus at 17dpr, showing loss of NeuroD1^+^ YFP^+^ cells in SOX4/11^iNestin^ mice. n= 3 animals. **(D)** Representative images showing loss of PROX1 (magenta) expression in recombined cells (green) in SOX4/11^iNestin^ mice 17dpr. Scale bar= 20µm **(E)** Quantification of PROX1^+^ YFP^+^ cells in the sub-granular zone of the dentate gyrus at 17dpr, showing loss of PROX1^+^ YFP^+^ cells in SOX4/11^iNestin^ mice. n= 3 animals. Data represented as mean ± SD; t test was used to determine significance.***P<0.001, **P<0.01, *P<0.05, ns = non-significant; dots represent individual animals

**Figure EV 3:**
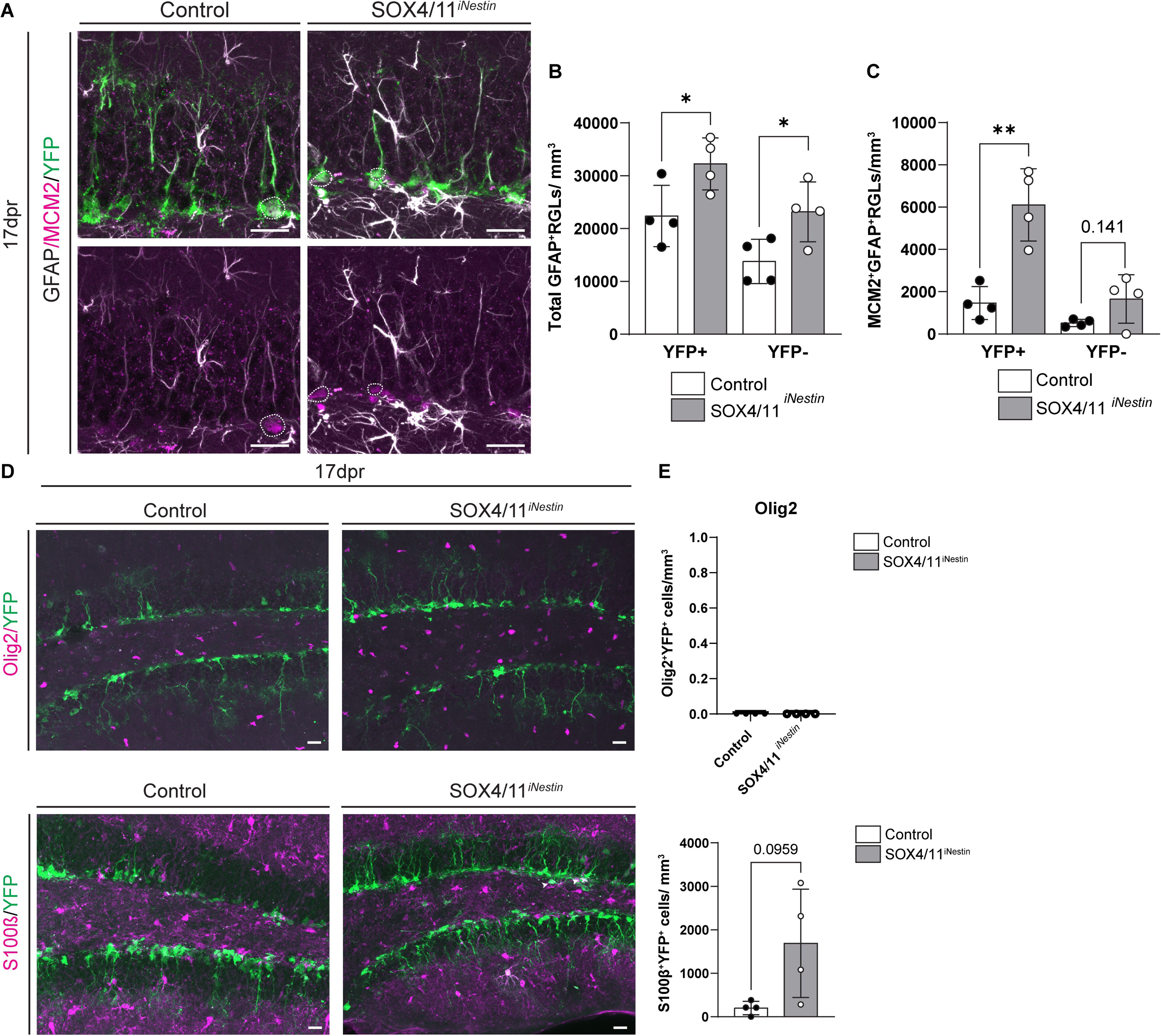
Data supporting Fig 3: **(A)** Representative images from control and SOX4/11^iNestin^ mice at 17dpr showing recombined (green) RGLs identified by GFAP expression (grey) and radial morphology in the subgranular zone. Activated RGLs (white circle) are identified by MCM2 expression (magenta). Scale bar = 20µm. **(B)** Quantification at 17dpr shows increase in recombined (YFP^+^) and non-recombined (YFP-) RGLs in SOX4/11^iNestin^ mice. n= 4 animals. **(C)** Quantification of activated RGLs at 17dpr shows a significant increase of recombined (YFP^+^) activated RGLs in SOX4/11^iNestin^ mice and a trend towards higher numbers of non-recombined (YFP-) activated RGLs in SOX4/11^iNestin^ mice. n= 4 animals. **(D)** Representative images showing Olig2 (magenta, upper panel) and S100β (magenta, lower panel) staining in the dentate gyrus of control and SOX4/11iNestin mice at 17dpr. Recombined cells (green). Scale bar = 20µm. **(E)** Quantification of Olig2^+^ and S100β^+^ cells at 17dpr. n= 4 animals. Data represented as mean ± SD; t test was used to determine significance.

**Figure EV 4:**
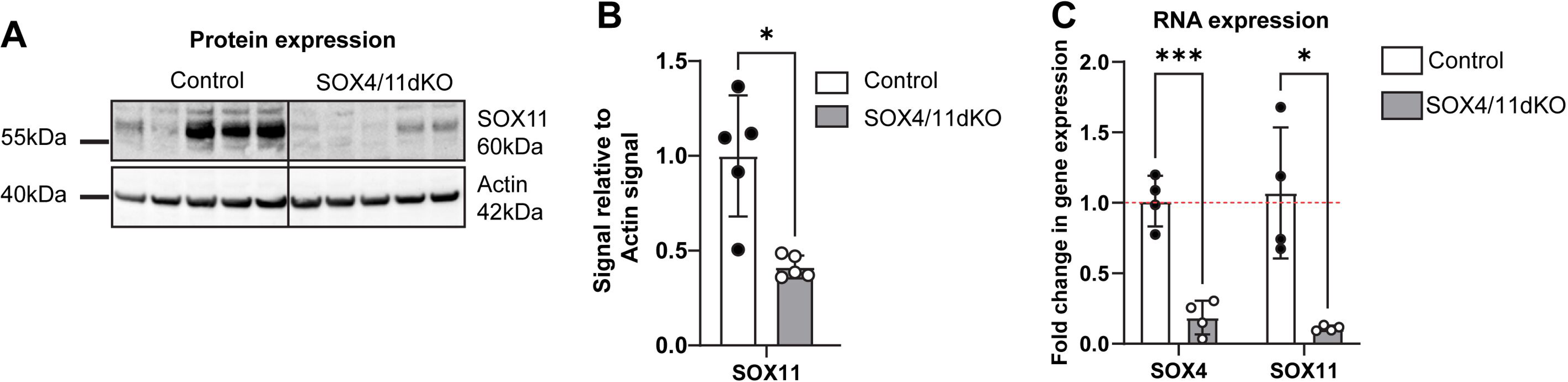
Data supporting Fig 4: **(A)** Immunoblot image for antibody against SOX11 showing the reduction in protein expression of SOX11 in the whole cells lysate 3 days post transduction with CAG-GFP and CAG-GFP-IRES-Cre MML-retroviruses. **(B)** Quantification from the western blot analysis showed a decrease in SOX11 expression relative to actin expression. n= 5 biological replicates. **(C)** Decrease in RNA expression of SOX4 and SOX11 quantified relative to housekeeping genes (hprt and RPL27). n=4 biological replicates. Statistical significance was calculated using t-test and quantification represented as mean ± SD. ***P<0.001, **P<0.01, *P<0.05, ns = non-significant.

**Figure EV 5:**
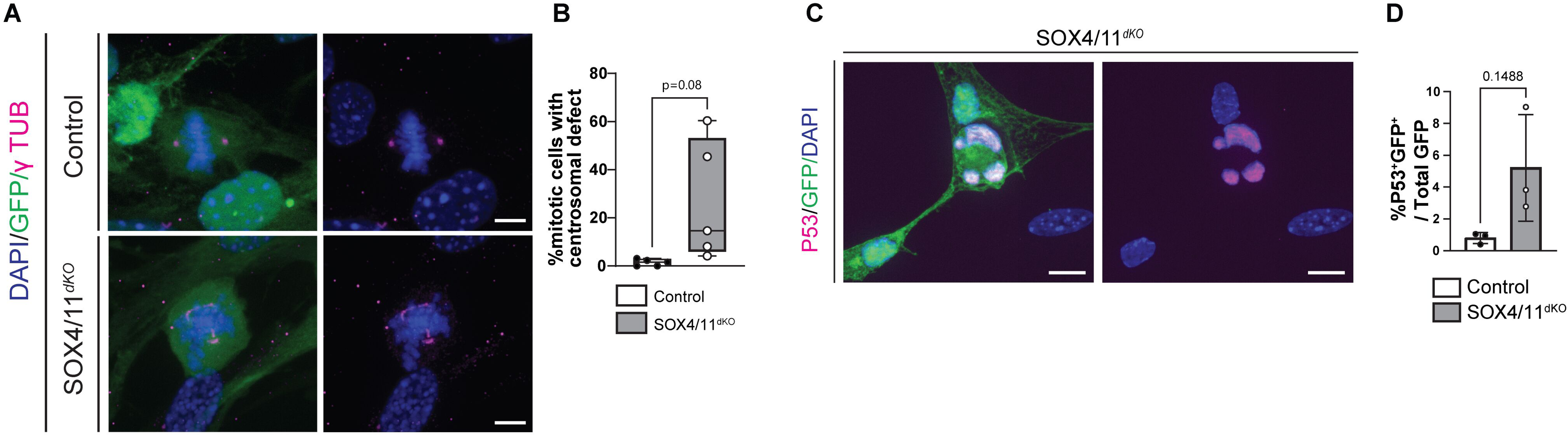
Data supporting Fig 5: **(A)** γ-Tubulin (γ-Tub, magenta) immunostaining of control and SOX4/11^dKO^ cells shows presence of multiple centrosome. Note the difference in morphology of the γ-Tub signal. γ-Tub signal in SOX4/11dKO is elongated rather than globular. Scale bar = 5µm **(B)** Percentage of transduced cells showing presence of centrosomal defects. At least 50 transduced metaphase were analyzed per condition. n=5 biological replicates. **(C)** Representative images showing presence of P53 (magenta) in transduced (green) multinucleated SOX4/11dKO cells. Scale bar = 10µm. **(D)** Quantification of percentage of transduced cells that showed P53 expression. n=3 biological replicates. Statistical significance was calculated using t-test and quantification represented as mean ± SD. ***P<0.001, **P<0.01, *P<0.05, ns = non-significant.

**Video EV5.1: live cell imaging of mitosis of control aNSPCs**

**Video EV5.2: live cell imaging of mitosis of SOX4/11dKO aNSPCs**

**Figure EV 7:**
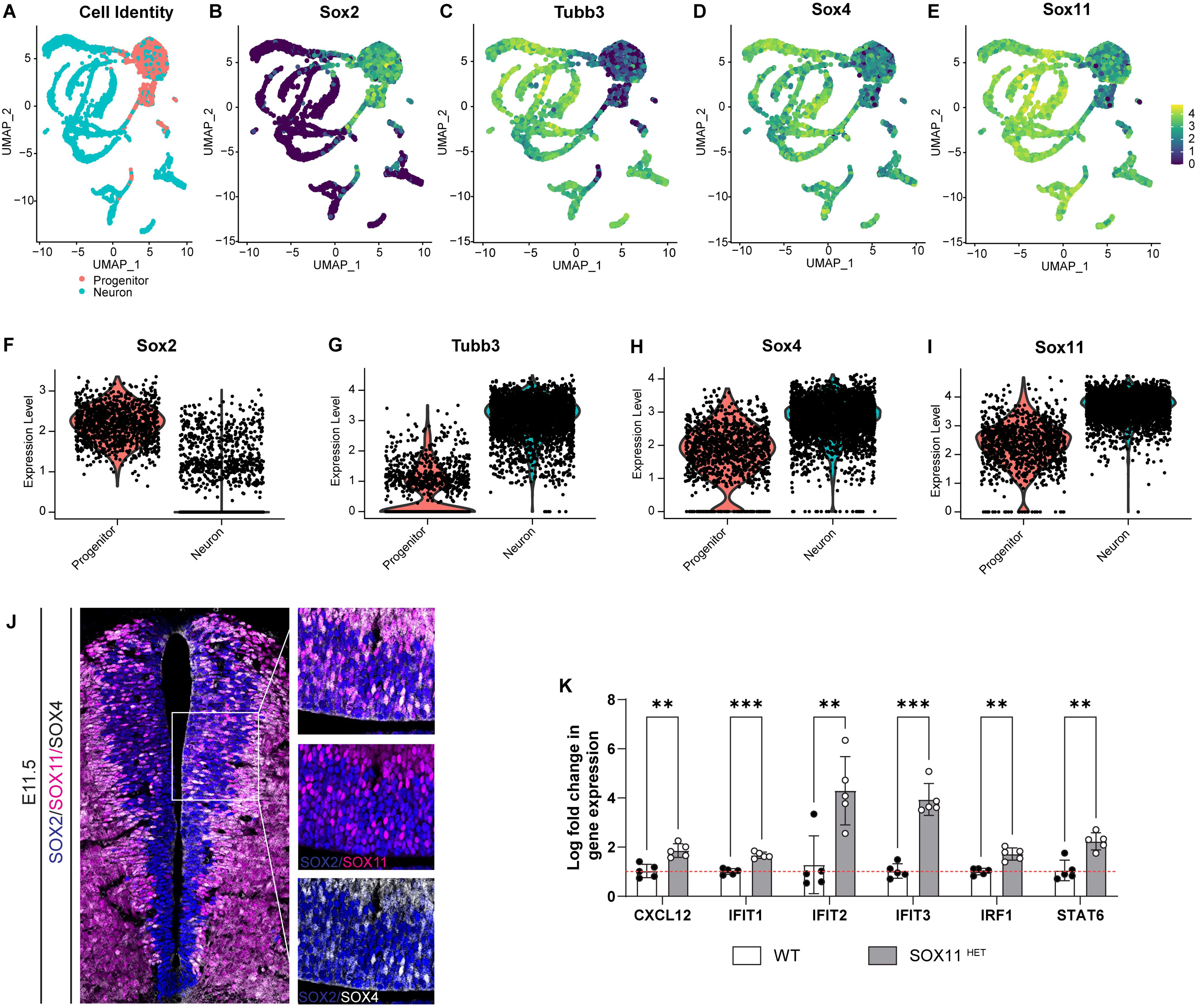
Data supporting Fig 7: **(A)** Clusters for Neurons and neural progenitors from ScRNA Seq datasets of E11.5 spinal cord [Dataref: (Delile et al., 2019)]. **(B-E)** UMAPs showing progenitor clusters enriched in SOX2 and neuronal clusters enriched for Tubb3 also expressing SOX4 and SOX11 mRNA. **(F-I)** Expression levels of SOX4 and SOX11 quantified in the SOX2 expressing progenitor and Tubb3 expressing neuronal population showing the expressing of SOX4 and SOX11 in the E11.5 spinal cord progenitors. **(J)** Immunohistochemical staining showing the Co-expression of SOX4 (blue) and SOX11 (magenta) protein with SOX2 (grey) expressing progenitors in the ventricular zone of E11.5 wildtype murine spinal cord. **(K)** qRT-PCR analysis from day 30 WT^Crispr^ and SOX11^Het^ organoids showing increased expression of inflammatory response molecules. n= 3 organoids from 2 batches. Statistical significance was calculated using t-test and quantification represented as mean ± SD. ***P<0.001, **P<0.01, *P<0.05, ns = non-significant.

## References

Abrous DN, Wojtowicz JM (2015) Interaction between Neurogenesis and Hippocampal Memory System: New Vistas. Cold Spring Harb Perspect Biol 7

Ahlenius H, Visan V, Kokaia M, Lindvall O, Kokaia Z (2009) Neural stem and progenitor cells retain their potential for proliferation and differentiation into functional neurons despite lower number in aged brain. J Neurosci 29: 4408–4419

Al-Jawahiri R, Foroutan A, Kerkhof J, McConkey H, Levy M, Haghshenas S, Rooney K, Turner J, Shears D, Holder M et al (2022) SOX11 variants cause a neurodevelopmental disorder with infrequent ocular malformations and hypogonadotropic hypogonadism and with distinct DNA methylation profile. Genet Med 24: 1261–1273

Arai Y, Pulvers JN, Haffner C, Schilling B, Nusslein I, Calegari F, Huttner WB (2011) Neural stem and progenitor cells shorten S-phase on commitment to neuron production. Nat Commun 2: 154

Bergsland M, Werme M, Malewicz M, Perlmann T, Muhr J (2006) The establishment of neuronal properties is controlled by Sox4 and Sox11. Genes Dev 20: 3475–3486

Bhattaram P, Penzo-Mendez A, Sock E, Colmenares C, Kaneko KJ, Vassilev A, Depamphilis ML, Wegner M, Lefebvre V (2010) Organogenesis relies on SoxC transcription factors for the survival of neural and mesenchymal progenitors. Nat Commun 1: 9

Boldrini M, Fulmore CA, Tartt AN, Simeon LR, Pavlova I, Poposka V, Rosoklija GB, Stankov A, Arango V, Dwork AJ et al (2018) Human Hippocampal Neurogenesis Persists throughout Aging. Cell Stem Cell 22: 589–599 e585

Breuss M, Fritz T, Gstrein T, Chan K, Ushakova L, Yu N, Vonberg FW, Werner B, Elling U, Keays DA (2016) Mutations in the murine homologue of TUBB5 cause microcephaly by perturbing cell cycle progression and inducing p53-associated apoptosis. Development 143: 1126–1133

Chen C, Lee GA, Pourmorady A, Sock E, Donoghue MJ (2015) Orchestration of Neuronal Differentiation and Progenitor Pool Expansion in the Developing Cortex by SoxC Genes. J Neurosci 35: 10629–10642

Contadini C, Monteonofrio L, Virdia I, Prodosmo A, Valente D, Chessa L, Musio A, Fava LL, Rinaldo C, Di Rocco G, Soddu S (2019) p53 mitotic centrosome localization preserves centrosome integrity and works as sensor for the mitotic surveillance pathway. Cell Death Dis 10: 850

Da Silva F, Zhang K, Pinson A, Fatti E, Wilsch-Brauninger M, Herbst J, Vidal V, Schedl A, Huttner WB, Niehrs C (2021) Mitotic WNT signalling orchestrates neurogenesis in the developing neocortex. EMBO J 40: e108041

Delile J, Rayon T, Melchionda M, Edwards A, Briscoe J, Sagner A (2019) Single cell transcriptomics reveals spatial and temporal dynamics of gene expression in the developing mouse spinal cord. Development 146

Demars M, Hu YS, Gadadhar A, Lazarov O (2010) Impaired neurogenesis is an early event in the etiology of familial Alzheimer’s disease in transgenic mice. J Neurosci Res 88: 2103–2117

Dumitru I, Paterlini M, Zamboni M, Ziegenhain C, Giatrellis S, Saghaleyni R, Björklund Å, Alkass K, Tata M, Druid H et al (2025) Identification of proliferating neural progenitors in the adult human hippocampus. Science 389: 58–63

Eriksson Ps, Perfilieva E, Björk-Eriksson T, Alborn A-M, Nordborg C, Peterson Da, & Gage FH (1998) Neurogenesis in the adult human hippocampus. NATURE MEDICINE 4

Fischer M, Schade AE, Branigan TB, Muller GA, DeCaprio JA (2022) Coordinating gene expression during the cell cycle. Trends Biochem Sci 47: 1009–1022

Flynn PJ, Koch PD, Mitchison TJ (2021) Chromatin bridges, not micronuclei, activate cGAS after drug-induced mitotic errors in human cells. Proc Natl Acad Sci U S A 118

Giandomenico SL, Mierau SB, Gibbons GM, Wenger LMD, Masullo L, Sit T, Sutcliffe M, Boulanger J, Tripodi M, Derivery E et al (2019) Cerebral organoids at the air-liquid interface generate diverse nerve tracts with functional output. Nat Neurosci 22: 669–679

Goncalves JT, Schafer ST, Gage FH (2016) Adult Neurogenesis in the Hippocampus: From Stem Cells to Behavior. Cell 167: 897–914

Hempel A, Pagnamenta AT, Blyth M, Mansour S, McConnell V, Kou I, Ikegawa S, Tsurusaki Y, Matsumoto N, Lo-Castro A et al (2016) Deletions and de novo mutations of SOX11 are associated with a neurodevelopmental disorder with features of Coffin-Siris syndrome. J Med Genet 53: 152–162

Hill AS, Sahay A, Hen R (2015) Increasing Adult Hippocampal Neurogenesis is Sufficient to Reduce Anxiety and Depression-Like Behaviors. Neuropsychopharmacology 40: 2368–2378

Hodge RD, Nelson BR, Kahoud RJ, Yang R, Mussar KE, Reiner SL, Hevner RF (2012) Tbr2 is essential for hippocampal lineage progression from neural stem cells to intermediate progenitors and neurons. J Neurosci 32: 6275–6287

Hoser M, Potzner MR, Koch JM, Bosl MR, Wegner M, Sock E (2008) Sox12 deletion in the mouse reveals nonreciprocal redundancy with the related Sox4 and Sox11 transcription factors. Mol Cell Biol 28: 4675–4687

Hoshiba Y, Toda T, Ebisu H, Wakimoto M, Yanagi S, Kawasaki H (2016) Sox11 Balances Dendritic Morphogenesis with Neuronal Migration in the Developing Cerebral Cortex. J Neurosci 36: 5775–5784

Imayoshi I, Sakamoto M, Ohtsuka T, Kageyama R (2009) Continuous neurogenesis in the adult brain. Dev Growth Differ 51: 379–386

Kempermann G, Gast D, Kronenberg G, Yamaguchi M, Gage FH (2003) Early determination and long-term persistence of adult-generated new neurons in the hippocampus of mice. Development 130: 391–399

Kempermann G, Song H, Gage FH (2015) Neurogenesis in the Adult Hippocampus. Cold Spring Harb Perspect Biol 7: a018812

Kletter T, Reusch S, Cavazza T, Dempewolf N, Tischer C, Reber S (2022) Volumetric morphometry reveals spindle width as the best predictor of mammalian spindle scaling. J Cell Biol 221

Kuhn HG, Toda T, Gage FH (2018) Adult Hippocampal Neurogenesis: A Coming-of-Age Story. J Neurosci 38: 10401–10410

Lancaster MA, Renner M, Martin CA, Wenzel D, Bicknell LS, Hurles ME, Homfray T, Penninger JM, Jackson AP, Knoblich JA (2013) Cerebral organoids model human brain development and microcephaly. Nature 501: 373–379

Lange C, Huttner WB, Calegari F (2009) Cdk4/cyclinD1 overexpression in neural stem cells shortens G1, delays neurogenesis, and promotes the generation and expansion of basal progenitors. Cell Stem Cell 5: 320–331

Lefebvre V, Dumitriu B, Penzo-Mendez A, Han Y, Pallavi B (2007) Control of cell fate and differentiation by Sry-related high-mobility-group box (Sox) transcription factors. Int J Biochem Cell Biol 39: 2195–2214

Levine MS, Holland AJ (2018) The impact of mitotic errors on cell proliferation and tumorigenesis. Genes Dev 32: 620–638

Mackenzie KJ, Carroll P, Martin CA, Murina O, Fluteau A, Simpson DJ, Olova N, Sutcliffe H, Rainger JK, Leitch A et al (2017) cGAS surveillance of micronuclei links genome instability to innate immunity. Nature 548: 461–465

McKinnon PJ (2013) Maintaining genome stability in the nervous system. Nat Neurosci 16: 1523–1529

Mitchell-Dick A, Chalem A, Pilaz LJ, Silver DL (2019) Acute Lengthening of Progenitor Mitosis Influences Progeny Fate during Cortical Development in vivo. Dev Neurosci 41: 300–317

Moreno-Jimenez EP, Flor-Garcia M, Terreros-Roncal J, Rabano A, Cafini F, Pallas-Bazarra N, Avila J, Llorens-Martin M (2019) Adult hippocampal neurogenesis is abundant in neurologically healthy subjects and drops sharply in patients with Alzheimer’s disease. Nat Med 25: 554–560

Mori T, Tanaka K, Buffo A, Wurst W, Kuhn R, Gotz M (2006) Inducible gene deletion in astroglia and radial glia--a valuable tool for functional and lineage analysis. Glia 54: 21–34

Mu L, Berti L, Masserdotti G, Covic M, Michaelidis TM, Doberauer K, Merz K, Rehfeld F, Haslinger A, Wegner M et al (2012) SoxC transcription factors are required for neuronal differentiation in adult hippocampal neurogenesis. J Neurosci 32: 3067–3080

Orth JD, Loewer A, Lahav G, Mitchison TJ (2012) Prolonged mitotic arrest triggers partial activation of apoptosis, resulting in DNA damage and p53 induction. Mol Biol Cell 23: 567–576

Phan TP, Holland AJ (2021) Time is of the essence: the molecular mechanisms of primary microcephaly. Genes Dev 35: 1551–1578

Pilaz LJ, McMahon JJ, Miller EE, Lennox AL, Suzuki A, Salmon E, Silver DL (2016) Prolonged Mitosis of Neural Progenitors Alters Cell Fate in the Developing Brain. Neuron 89: 83–99

Pilz G-A, Bottes S, Betizeau M, Jörg DJ, Carta S, April S, Simons BD, Helmchen F, Jessberger S (2018) Live imaging of neurogenesis in the adult mouse hippocampus. Science 359: 658–662

Rasetto NB, Giacomini D, Berardino AA, Waichman TV, Beckel MS, Di Bella DJ, Brown J, Davies--Sala MG, Gerhardinger C, Lie DC et al (2024) Transcriptional dynamics orchestrating the development and integration of neurons born in the adult hippocampus. Science Advances

Rehen SK, McConnell MJ, Kaushal D, Kingsbury MA, Yang AH, Chun J (2001) Chromosomal variation in neurons of the developing and adult mammalian nervous system. PNAS 98: 13361– 13366

Rigato C, Buckinx R, Le-Corronc H, Rigo JM, Legendre P (2011) Pattern of invasion of the embryonic mouse spinal cord by microglial cells at the time of the onset of functional neuronal networks. Glia 59: 675–695

Sazonova EV, Petrichuk SV, Kopeina GS, Zhivotovsky B (2021) A link between mitotic defects and mitotic catastrophe: detection and cell fate. Biol Direct 16: 25

Schaffner I, Minakaki G, Khan MA, Balta EA, Schlotzer-Schrehardt U, Schwarz TJ, Beckervordersandforth R, Winner B, Webb AE, DePinho RA et al (2018) FoxO Function Is Essential for Maintenance of Autophagic Flux and Neuronal Morphogenesis in Adult Neurogenesis. Neuron 99: 1188–1203 e1186

Schaffner I, Wittmann MT, Vogel T, Lie DC (2023) Differential vulnerability of adult neurogenic niches to dosage of the neurodevelopmental-disorder linked gene Foxg1. Mol Psychiatry 28: 497–514

Schincariol-Manhe B, Campagnolo E, Spineli-Silva S, de Leeuw N, Correia-Costa GR, Pessoa A, de Souza CFM, Stevens C, Javaher P, Scallet HF et al (2024) Novel variants in the SOX11 gene: clinical description of seven new patients. Eur J Hum Genet 32: 1640–1646

Shi L, Qalieh A, Lam MM, Keil JM, Kwan KY (2019) Robust elimination of genome-damaged cells safeguards against brain somatic aneuploidy following Knl1 deletion. Nat Commun 10: 2588

Shin J, Berg DA, Zhu Y, Shin JY, Song J, Bonaguidi MA, Enikolopov G, Nauen DW, Christian KM, Ming GL, Song H (2015) Single-Cell RNA-Seq with Waterfall Reveals Molecular Cascades underlying Adult Neurogenesis. Cell Stem Cell 17: 360–372

Sierra A, Encinas JM, Deudero JJ, Chancey JH, Enikolopov G, Overstreet-Wadiche LS, Tsirka SE, Maletic-Savatic M (2010) Microglia shape adult hippocampal neurogenesis through apoptosis-coupled phagocytosis. Cell Stem Cell 7: 483–495

Spalding KL, Bergmann O, Alkass K, Bernard S, Salehpour M, Huttner HB, Bostrom E, Westerlund I, Vial C, Buchholz BA et al (2013) Dynamics of hippocampal neurogenesis in adult humans. Cell 153: 1219–1227

Steib K, Schaffner I, Jagasia R, Ebert B, Lie DC (2014) Mitochondria modify exercise-induced development of stem cell-derived neurons in the adult brain. J Neurosci 34: 6624–6633

Takaki T, Millar R, Hiley CT, Boulton SJ (2024) Micronuclei induced by radiation, replication stress, or chromosome segregation errors do not activate cGAS-STING. Mol Cell 84: 2203–2213 e2205

Tashiro A, Zhao C, Gage FH (2006) Retrovirus-mediated single-cell gene knockout technique in adult newborn neurons in vivo. Nat Protoc 1: 3049–3055

Terreros-Roncal J, Moreno-Jiménez EP, Flor-García M, Rodríguez-Moreno CB, Trinchero MF, Cafini F, Rábano A, Llorens-Martín M (2021) Impact of neurodegenerative diseases on human adult hippocampal neurogenesis. Science 374: 1106–1113

Thein DC, Thalhammer JM, Hartwig AC, Crenshaw EB, 3rd, Lefebvre V, Wegner M, Sock E (2010) The closely related transcription factors Sox4 and Sox11 function as survival factors during spinal cord development. J Neurochem 115: 131–141

Tobin MK, Musaraca K, Disouky A, Shetti A, Bheri A, Honer WG, Kim N, Dawe RJ, Bennett DA, Arfanakis K, Lazarov O (2019) Human Hippocampal Neurogenesis Persists in Aged Adults and Alzheimer’s Disease Patients. Cell Stem Cell 24: 974–982 e973

Tsurusaki Y, Koshimizu E, Ohashi H, Phadke S, Kou I, Shiina M, Suzuki T, Okamoto N, Imamura S, Yamashita M et al (2014) De novo SOX11 mutations cause Coffin-Siris syndrome. Nat Commun 5: 4011

Tuncdemir SN, Lacefield CO, Hen R (2019) Contributions of adult neurogenesis to dentate gyrus network activity and computations. Behav Brain Res 374: 112112

Turan S, Boerstler T, Kavyanifar A, Loskarn S, Reis A, Winner B, Lie DC (2019) A novel human stem cell model for Coffin-Siris syndrome-like syndrome reveals the importance of SOX11 dosage for neuronal differentiation and survival. Hum Mol Genet 28: 2589–2599

Unger MS, Marschallinger J, Kaindl J, Hofling C, Rossner S, Heneka MT, Van der Linden A, Aigner L (2016) Early Changes in Hippocampal Neurogenesis in Transgenic Mouse Models for Alzheimer’s Disease. Mol Neurobiol 53: 5796–5806

Wang Y, Lin L, Lai H, Parada LF, Lei L (2013) Transcription factor Sox11 is essential for both embryonic and adult neurogenesis. Dev Dyn 242: 638–653

Wegner M (2011) SOX after SOX: SOXession regulates neurogenesis. Genes Dev 25: 2423–2428

Yang AH, Kaushal D, Rehen SK, Kriedt K, Kingsbury MA, McConnell MJa, Chun J (2003) Chromosome Segregation Defects Contribute to Aneuploidy in Normal Neural Progenitor Cells. The Journal of Neuroscience 23: 10454 –10462

Zhou Y, Su Y, Li S, Kennedy BC, Zhang DY, Bond AM, Sun Y, Jacob F, Lu L, Hu P et al (2022) Molecular landscapes of human hippocampal immature neurons across lifespan. Nature 607: 527–533

